# Master clock‒thalamic‒prefrontal circuit controls circadian social priority

**DOI:** 10.64898/2025.12.05.691743

**Authors:** Jihoon Kim, Jinseon Yu, Boil Kim, Inah Park, Minseok Kim, Jimin Lee, Mijung Choi, Cheol Song, Yulong Li, Kyungjin Kim, Jong-Cheol Rah, Han Kyoung Choe

## Abstract

Social behaviors arise from complex brain operations that adapt to changing social contexts and internal states. However, how internal states guide the resolution of competing social goals remains unclear. Here, we show that prioritization between competing instincts follows a circadian pattern via a circuit linking the suprachiasmatic nucleus (SCN) to the medial prefrontal cortex (mPFC). Specifically, SCN^VIP^ neurons project to Vipr2-expressing neurons in the nucleus reuniens (RE), which innervate the mPFC to regulate social prioritization. Vipr2 signaling in the RE is essential for circadian modulation of appetitive and consummatory social behaviors, aligning social behaviors with clock-regulated reproductive system. Overall, this SCN-RE-mPFC circuit provides a neural principle through which the master clock coordinates the prefrontal cortex to resolve a conflict of motivated behaviors by referencing an internal state.

## Introduction

Social behavior encompasses a diverse repertoire of behaviors, including mating, attacking, nurturing, and investigating, each serving distinct purposes (*1–3*). In complex social situations, organisms are simultaneously exposed to multiple social cues that evoke competing motivations, necessitating effective mechanisms for behavioral prioritization (*4–6*). Although the circuitry underlying many innate social behaviors has been extensively studied, the neural mechanism responsible for such behavioral prioritization between different behavioral expressions remains elusive.

A potential mechanism for social behavioral prioritization is circadian rhythm. It plays a crucial role in regulating daily homeostasis and maintaining biological rhythms (*7–9*). Numerous physiological and neural processes exhibit 24-hr rhythmicity governed by the suprachiasmatic nucleus (SCN), a hypothalamic region known as the master clock (*10–12*). SCN neurons maintain synchronized molecular clockworks, enabling self-sustaining circadian rhythm independent of external light signals, and coordinate circadian rhythm across the brain via neural and humoral pathways (*13–15*). Recent studies demonstrated that many behaviors display SCN-driven circadian rhythmicity (*16–20*), resulting in time-dependent expression of distinct behavioral outcomes. However, although several social behaviors display circadian rhythmicity, whether circadian rhythm prioritizes social behaviors in a time-dependent manner remains unknown. This study aimed to investigate the influence of circadian rhythm on social behaviors by developing a behavioral paradigm to assess behavioral prioritization between two distinct social cues.

## Results

### Circadian rhythm in social priority

To evaluate prioritization between competing social goals, we developed a social priority test that drives behavioral choice by establishing an approach‒approach conflict between sexual preference and social novelty. The protocol comprises two stages: a familiarization session and a subsequent prioritization session (Fig. 1A). During familiarization, a male subject mouse was placed in a three-chamber arena containing a novel female and a non-social object in separate chambers. During prioritization, a familiarized female and a novel male were presented in a counterbalanced, unbiased manner. The test was conducted at four circadian time points (CT00, 06, 12, 18). During the familiarization session, male subject mice consistently spent more time exploring the female cage (fig. S1A, B). In the prioritization session, they explored both the familiar female cage and the novel male cage, with preferences varying by circadian time (Fig. 1B, C). At CT12, the onset of the active phase, social novelty preference dominated, whereas sexual preference was enhanced at CT18, the mid-active phase. The total distance travelled and overall interaction time did not differ across time points (Fig. 1D, E; and fig. S1C, D).

**Fig. 1.**
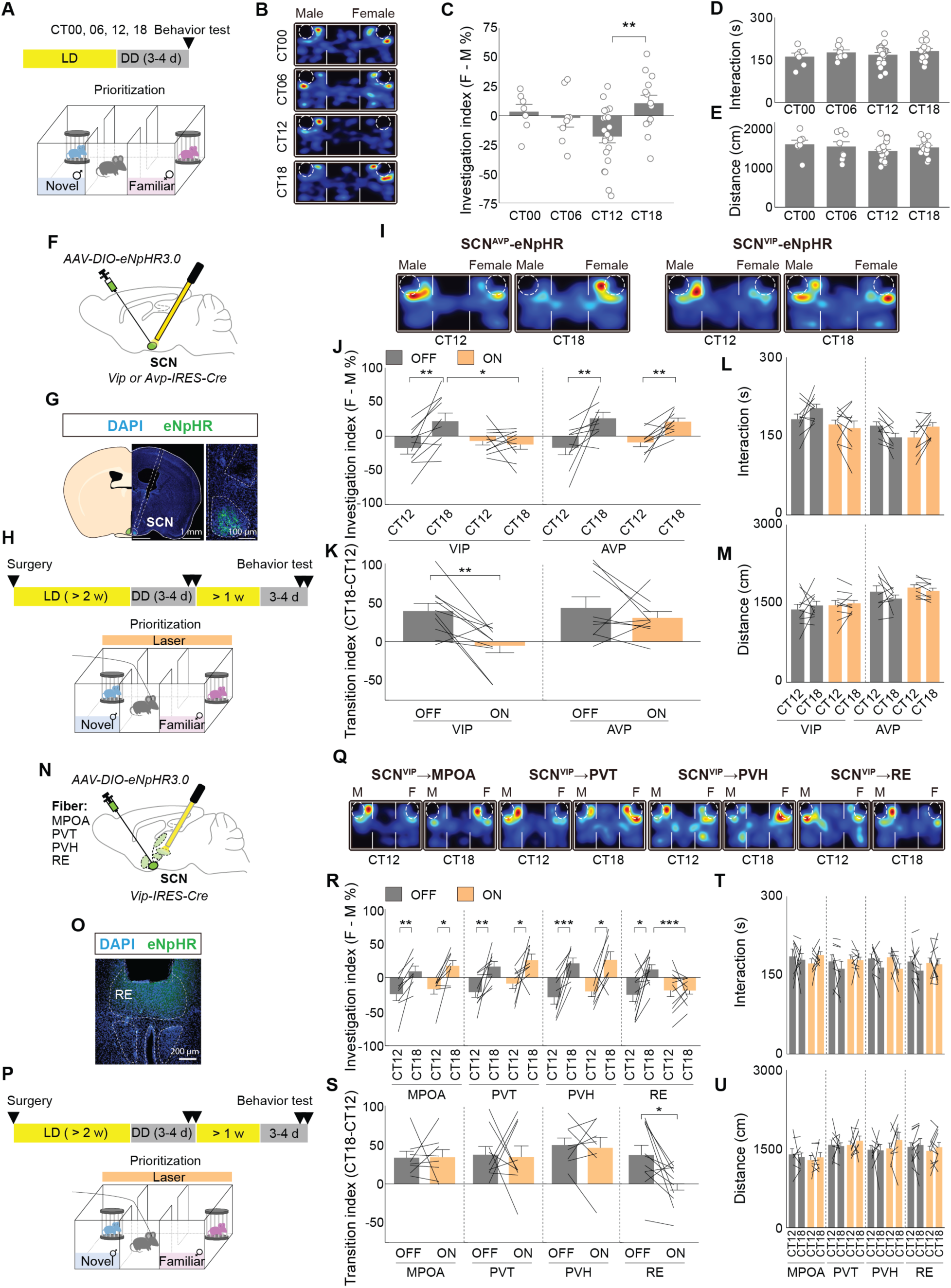
SCN^VIP^ projection to RE is required for circadian rhythm in social priority. (**A**) Experimental scheme of social priority assay. (**B**) Representative heatmap visualization of subject mice during social priority assay in each time point. (**C** to **E**) Quantification of investigation index (C), total interaction (D), and total distance moved (E) during social priority assay in each time point. (**F** to **G**) Schematics for viral delivery (F) and an example image showing viral expression and optic fiber placement (G). (**H**) Experimental scheme of social priority assay with optogenetic inhibition. (**I**) Representative heatmap visualization of subject mice during social priority assay with optogenetic inhibition in each time point. (**J** to **M**) Quantification of investigation index (J), transition index (K), total interaction (L), and total distance moved (M) during social priority assay with or without optogenetic inhibition. (**N-O**) Schematics for viral delivery (N) and an example image showing viral expression and optic fiber placement (O). (**P**) Experimental scheme of social priority assay with optogenetic inhibition. (**Q**) Representative heatmap visualization of subject mice during social priority assay with optogenetic inhibition in each time point. (**R** to **U**) Quantification of investigation index (R), transition index (S), total interaction (T), and total distance moved (U) during social priority assay with or without optogenetic inhibition. n = 7, 8, 21, 13 for CT00, 06, 12, 18 groups in (C to E); n = 10 for SCN^VIP^-eNpHR3.0 animals and n = 9 for SCN^AVP^-eNpHR3.0 animals in (J to M); n = 8 for SCN^VIP^-MPOA-eNpHR3.0, n = 9 for SCN^VIP^-PVT-eNpHR3.0, n = 8 for SCN^VIP^-PVH-eNpHR3.0, and n = 11 for SCN^VIP^-RE-eNpHR3.0 animals (R to U). For (C to E), One-way ANOVA test followed by Tukey’s HSD test; for (J), (L), (M), (R), (T), and (U), RM Two-way ANOVA test followed by pairwise paired t-test or non-parametric RM Two-way ANOVA test followed by Wilcoxon Signed-Rank test; for (K) and (S), paired t-test was performed. Each dot represents an individual animal. Each line represents a repeated experiment within individual animal. Data are presented as mean ± SEM. \*\*\**P* < 0.001; \*\**P* < 0.01; \**P* < 0.05.

We next investigated whether the individual social instincts underlying the approach‒approach conflict exhibit intrinsic circadian rhythmicity. Sexual preference remained constant across circadian times, with no changes in control metrics such as total interaction time or distance travelled (fig. S1E‒H). Similarly, social novelty, measured using the social recognition test, showed no circadian variance in preference, interaction time, and distance travelled (fig. S1I‒L). We further investigated whether female mice also exhibit a circadian rhythm of social priority. In female subjects, the social priority test revealed no significant time-of-day differences during either the familiarization or prioritization sessions, nor in total interaction time or distance travelled (fig. S2A‒D). These results were independent of estrus cycle stage. Likewise, neither the sexual preference test nor the social recognition test in females revealed circadian variances in preference, total interaction time, or locomotion (fig. S2H‒O).

A following question was whether the modulation of social priority originates from the internal time of the subject or the target. To disentangle these factors, we designed a misalignment experiment focusing on two pivotal time points, CT12 and CT18, where priority shifts from zero (fig. S3A, B). We created four experimental groups by mismatching subjects and targets across these two time points. During familiarization, all groups showed comparable investigation index and interaction time, with only minor variations in distance travelled (fig. S3C‒E). Conversely, a significant difference emerged during prioritization. Subjects tested at CT12 exhibited a novelty-seeking bias regardless of the target’s circadian time, whereas subjects tested at CT18 showed a consistent sexual preference bias (fig. S3F). Across these combinations, total interaction time did not differ, and although the difference in distance travelled was significant, the magnitude was marginal (fig. S3G, H).

### Master clock to the thalamic circuit for the circadian transition of social priority

Subsequently, we investigated how the circadian clock shapes daily patterns of social priority. Mammalian master clock resides in the SCN, which conveys circadian information to the brain and peripheral organs via neural and humoral pathways (*21*, *22*). The SCN comprises two major neural populations: vasoactive intestinal peptide–expressing (VIP[+]) and arginine vasopressin–expressing (AVP[+]) neurons. To determine which population mediates circadian control of social priority, we selectively inhibited SCN^VIP^ or SCN^AVP^ neurons. We injected AAV-DIO-eNpHR3.0 into the SCN of VIP-IRES-Cre or AVP-IRES-Cre mice and implanted optical fibers above the SCN (Fig. 1F, G; fig. S4A, B). After incubation and entrainment, the social priority tests were performed with optogenetic inhibition (Fig. 1H). During the familiarization session, optogenetic inhibition did not affect the investigation of female, total interaction, or locomotion (fig. S4C‒E). Conversely, inhibiting SCN^VIP^ neurons abolished the sexual preference bias at CT18, whereas inhibiting SCN^AVP^ neurons did not alter the transition of social priority between CT12 and CT18 (Fig. 1I‒K). Total interaction time and distance traveled during the prioritization session were unaffected (Fig. 1L, M). These results show that SCN^VIP^ neurons shape the circadian pattern of social priority.

To investigate the neural mechanism by which SCN^VIP^ neurons regulate circadian social prioritization, we optogenetically inhibited SCN^VIP^ neuronal terminals in 4 brain areas, including the medial preoptic area (MPOA), paraventricular thalamic nucleus (PVT), paraventricular hypothalamic nucleus (PVH), and nucleus reuniens (RE). Optical fibers were implanted above individual SCN^VIP^ target regions in SCN^VIP^-specific eNpHR3.0-expressing mice (Fig. 1N, O; fig. S5A‒H), and social priority was assessed during terminal inhibition (Fig. 1P). Optogenetic inhibition did not affect the investigation index, total interaction time, or distance travelled during the familiarization session (fig. S5I‒K). In the prioritization session, although inhibiting SCN^VIP^ projection targeting MPOA, PVT, or PVH did not alter social priority at CT12 and 18, SCN^VIP^-RE projection inhibition blocked the sexual preference bias at CT18 (Fig. 1Q, R) and eliminated the transition in social priority between CT12 and 18 (Fig. 1S). Across all groups, distance travelled during the prioritization session remained unaffected (Fig. 1T, U).

### VIP receptor 2 (VIPR2) signaling in the RE

Next, we examined whether VIP peptide signaling mediates the circadian pattern of social priority and, if so, through which receptor. VIP, a 28-amino-acid-long peptide, signals through two receptors, VIPR1 and VIPR2. Using intracerebroventricular injections, we pharmacologically blocked pan-VIPR, VIPR1, and VIPR2, respectively, and investigated social priority transitions (Fig. 2A‒C, and fig. S6A). Blocking both receptors abolished the sexual preference bias at CT18, an effect recapitulated by VIPR2 antagonism (Fig. 2D, E), whereas VIPR1 inhibition has no behavioral effect. This pattern is obvious in the transition index (Fig. 2F). Although total interaction time during the prioritization session was comparable to vehicle-treated controls, locomotor activity was affected by VIPR antagonist infusion (Fig. 2G, H). During familiarization, female investigation of animals infused with pan-VIPR antagonist at CT12 was significantly higher; however, most behaviors were comparable to the vehicle-injected group (fig. S6B‒D).

**Fig. 2.**
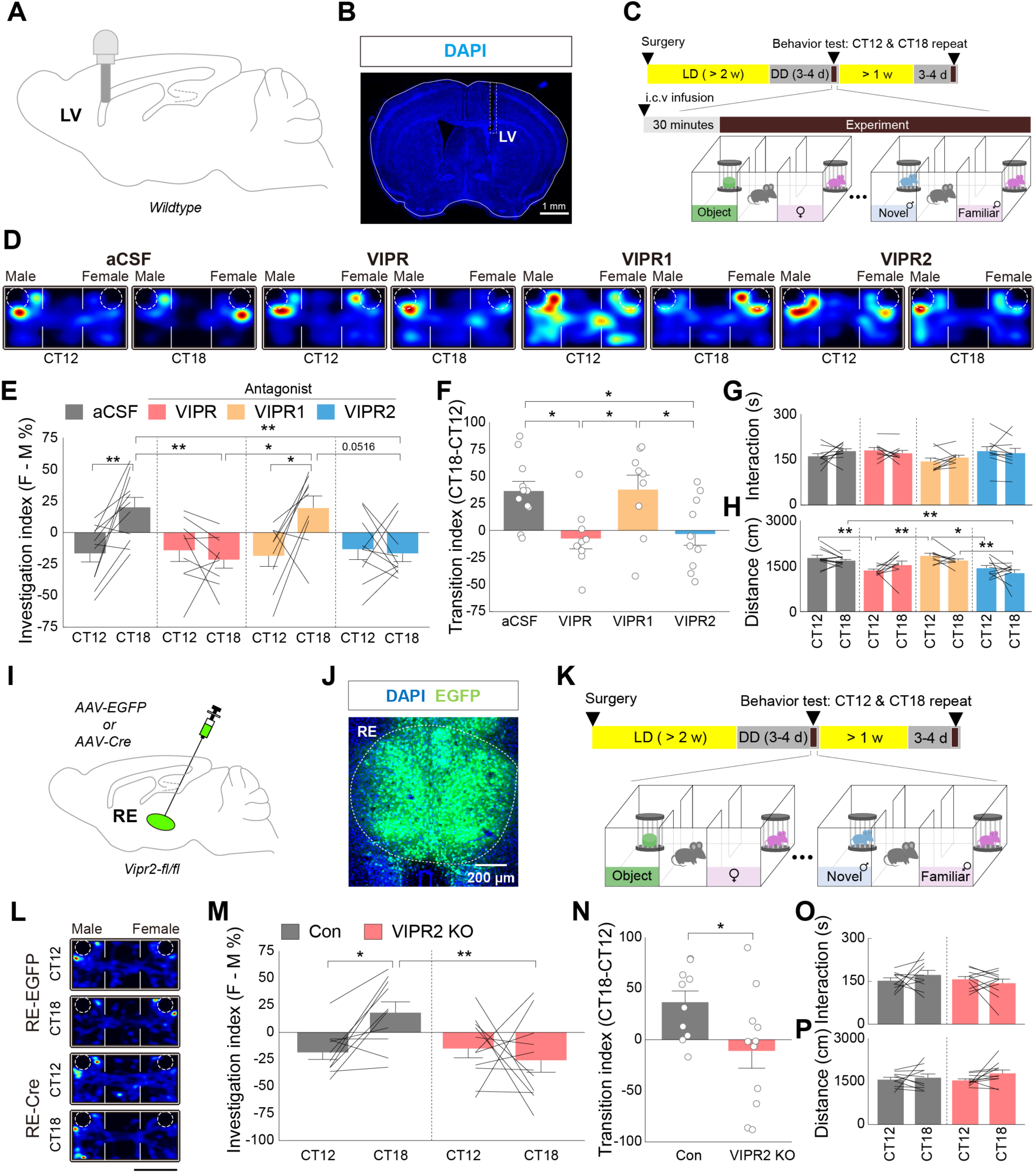
RE^VIPR2^ neuron mediates circadian rhythm in social priority. (**A** to **B**) Schematics for cannula implantation (A) and an example image showing cannula placement (B). (**C**) Experimental scheme of social priority assay with systematic infusion of various pharmacological agents for VIP signaling. (**D**) Representative heatmap visualization of subject mice during social priority assay with pharmacological inhibition. (**E** to **H**) Quantification of investigation index (E), transition index (F), total interaction (G), and total distance moved (H) during social priority assay with pharmacological inhibition. (**I** to **J**) Schematics for viral delivery (I) and an example image showing viral expression (J). (**K**) Experimental scheme of social priority assay with or without genetic ablation of Vipr2 in RE. (**L**) Representative heatmap visualization of subject mice during social priority assay with or without genetic ablation of Vipr2 in RE. (**M** to **P**) Quantification of investigation index (M), transition index (N), total interaction (O), and total distance moved (P) during social priority assay with or without genetic ablation of Vipr2 in RE. n = 11, 9, 9, 10 for aCSF, VIPR, VIPR1, and VIPR2 antagonist treatment groups in (E to H); n = 10, 11 for RE-EGFP and RE-Cre groups in (M to P). For (E), (G), (H), (M), (O), and (P), Two-way mixed ANOVA test followed by pairwise paired t-test and pairwise independent-sample t-test or non-parametric Two-way mixed ANOVA test followed by pairwise Wilcoxon Signed-Rank test and pairwise Wilcoxon Rank-Sum test; for (F), One-way ANOVA test followed by Tukey’s HSD test; for (N), unpaired t-test was performed. Each dot represents an individual animal. Each line represents a repeated experiment within individual animal. Data are presented as mean ± SEM. \*\**P* < 0.01; \**P* < 0.05.

To visualize the Vipr2-expressing neurons within the SCN^VIP^-RE circuit, we crossed Vipr2-IRES-Cre mice with Ai14 reporter mice, revealing tdTomato reporter expression in the RE, specifically within the same subregion targeted by SCN^VIP^ neuron projections (fig. S7A‒D). We then examined the function of VIPR2 signaling by conditionally knocking out Vipr2 in the RE (Fig. 2I‒K; fig. S7E, F). Compared with EGFP-expressing controls, RE-specific Vipr2 knockout (KO) significantly displayed a shift toward social novelty at CT18 (Fig. 2L, M), eliminating the normal transition between CT12 and 18 (Fig. 2N). Travelled distance and total interaction time were unchanged (Fig. 2O, P), as were behaviors during the familiarization session (fig. S7G‒I).

Behavioral effects of pharmacological and genetic Vipr2 manipulation were most prominent at CT12 and CT18, prompting us to examine whether VIP levels in the RE vary across these time points. We expressed a GPCR-based VIP sensor (GRAB_VIP1.0_) in the RE of Vipr2-IRES-Cre knock-in mice and measured sensor fluorescence at CT12 and CT18 (Fig. 3A‒C) (*23*). GRAB_VIP1.0_ intensities were higher at CT18 than at CT12 (Fig. 3D, E), indicating an increased RE VIPergic tone at CT18. We then examined whether this increase in VIP tone differentially modulates the neural activity of RE^Vipr2^ neurons in response to social cues. To confirm this, bulk calcium activity was recorded during the social priority assay at CT12 and CT18 (fig. S8A, B), and calcium transients were aligned to the onset of investigation of either a novel male or a familiar female. No cue-evoked calcium transients were detected (fig. S8C), and overall calcium activity did not differ between CT12 and CT18. Combined with GRAB_VIP1.0_ recording, these results suggest that RE^Vipr2^ neuronal activity does not directly encode social cue responsiveness.

**Fig. 3.**
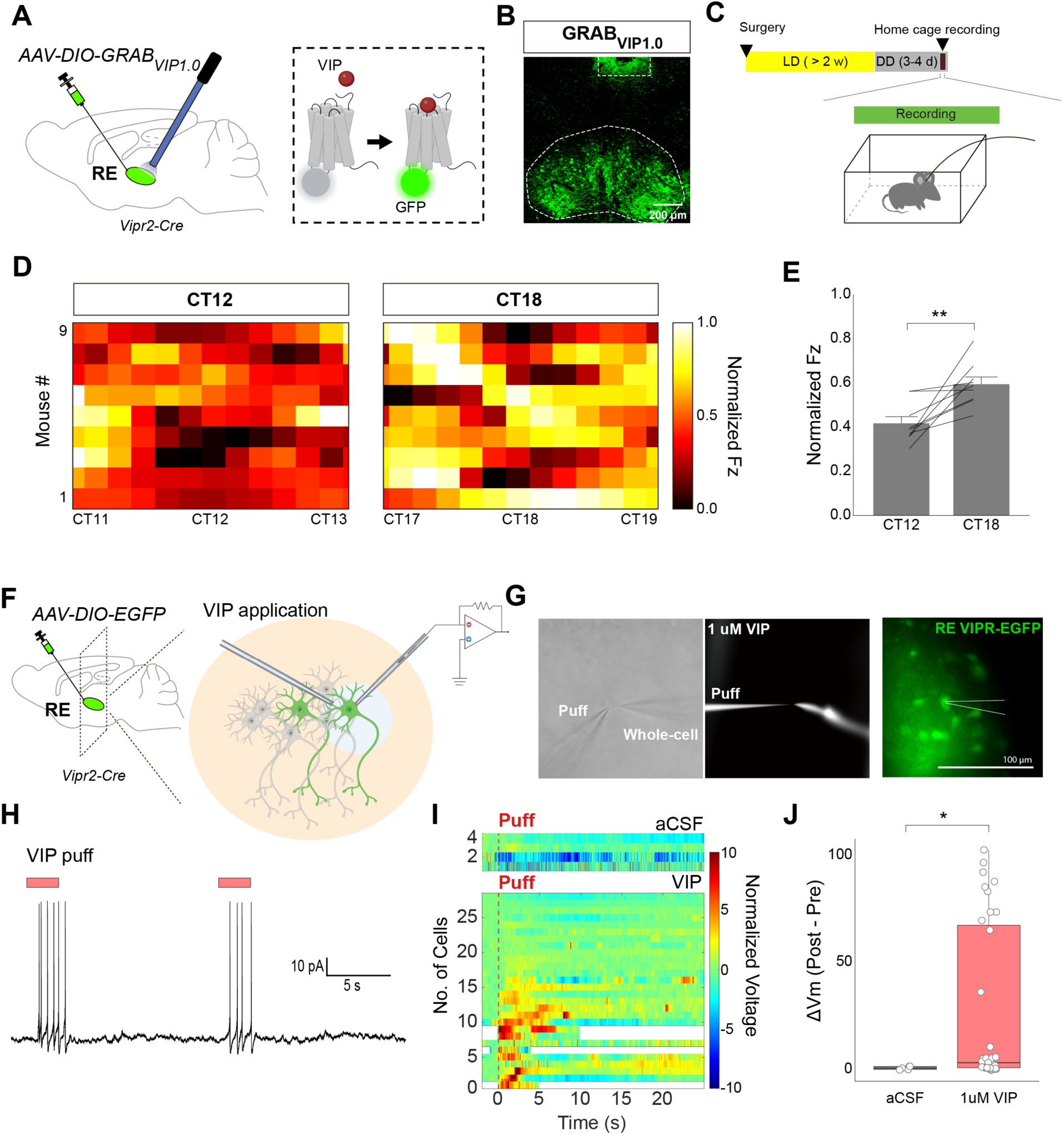
Circadian time-dependent fluctuation of VIPR signaling in RE^VIPR2^ neuron. (**A** to **B**) Schematics for viral delivery (A) and an example image showing viral expression and optic fiber placement (B). (**C**) Experimental scheme of VIPR activity recording in freely moving animals. (**D** to **E**) Heatmap visualization of VIPR activity (D) and quantification of normalized VIPR activity (E) across two circadian timepoints. (**F**) Schematics for viral delivery and acute slice section for electrophysiological recording. (**G**) An example image showing Vipr2-expressing neuron with puff pipette. (**H**) Representative current-clamp trace showing a depolarizing response evoked by VIP agonist puff. Black bar indicates the timing of puff application. (**I**) Heatmap visualization of firing rates computed in cells that exhibited spontaneous EPSPs. Firing rate was calculated using a 500-ms sliding window. Top: aCSF puff (control); Bottom: VIP puff. (**J**) The difference between the average Vm during the 2-second baseline period (Pre) and the peak depolarization following the drug infusion (Post) was compared between control and VIP-treated cells. n = 9 for GRAB_VIP1.0_ recording in (D to E); n = 5, 35 for aCSF and VIP infusion in (H to J). For (E), paired t-test; for (J), Wilcoxon Rank-Sum test were performed. Each line represents a repeated experiment within individual animal or cell. Data are presented as mean ± SEM. \*\**P* < 0.01; \**P* < 0.05. Panel **A** (right) and **F** (right) were created with BioRender.com.

Next, we investigated the electrophysiological effect of VIPR2 activation in RE^Vipr2^ neurons via whole-cell patch-clamp recordings (Fig. 3F). VIP peptide puffs were delivered near fluorescently labeled RE^Vipr2^ neurons through AAV-DIO-EGFP expression (Fig. 3G), resulting in robust VIP peptide puff-induced depolarizing voltage responses (Fig. 3H). In responsive neurons, a VIP puff application elicited rapid depolarization, with minimal residual current following washout. VIP puff significantly depolarized RE^Vipr2^ neurons from the baseline with varying degree (Fig. 3I, J). Together, these findings indicate that VIPergic tone in the RE is elevated at CT18, and that elevated VIP signaling electrophysiologically activates RE^Vipr2^ neurons.

### Thalamo-cortical circuit for social priority

Through extensive connectivity with various brain areas, including the medial prefrontal cortex (mPFC) and hippocampus, RE has been implicated in social behavior and flexible behavioral control, making it a strong candidate for regulating social priority(*24–27*). To determine the causal role of RE^Vipr2^ neurons in prioritizing between social novelty and sexual preference, we bidirectionally manipulated their activity at CT12 and CT18 using optogenetics (Fig. 4A‒C; fig. S9A, B). Optogenetic inhibition through eNpHR consistently biased mice toward investigating the novel male at both time points (Fig. 4D, E). Conversely, optogenetic activation via ChR2 biased investigation toward the familiar female, enhancing sexual preference. Crucially, both manipulations abolished the normal circadian transition in social priority (Fig. 4F). These effects were specific, as neither total interaction time nor locomotor activity was altered (Fig. 4G, H), and only minor changes were observed during the familiarization session (fig. S9C‒E).

**Fig. 4.**
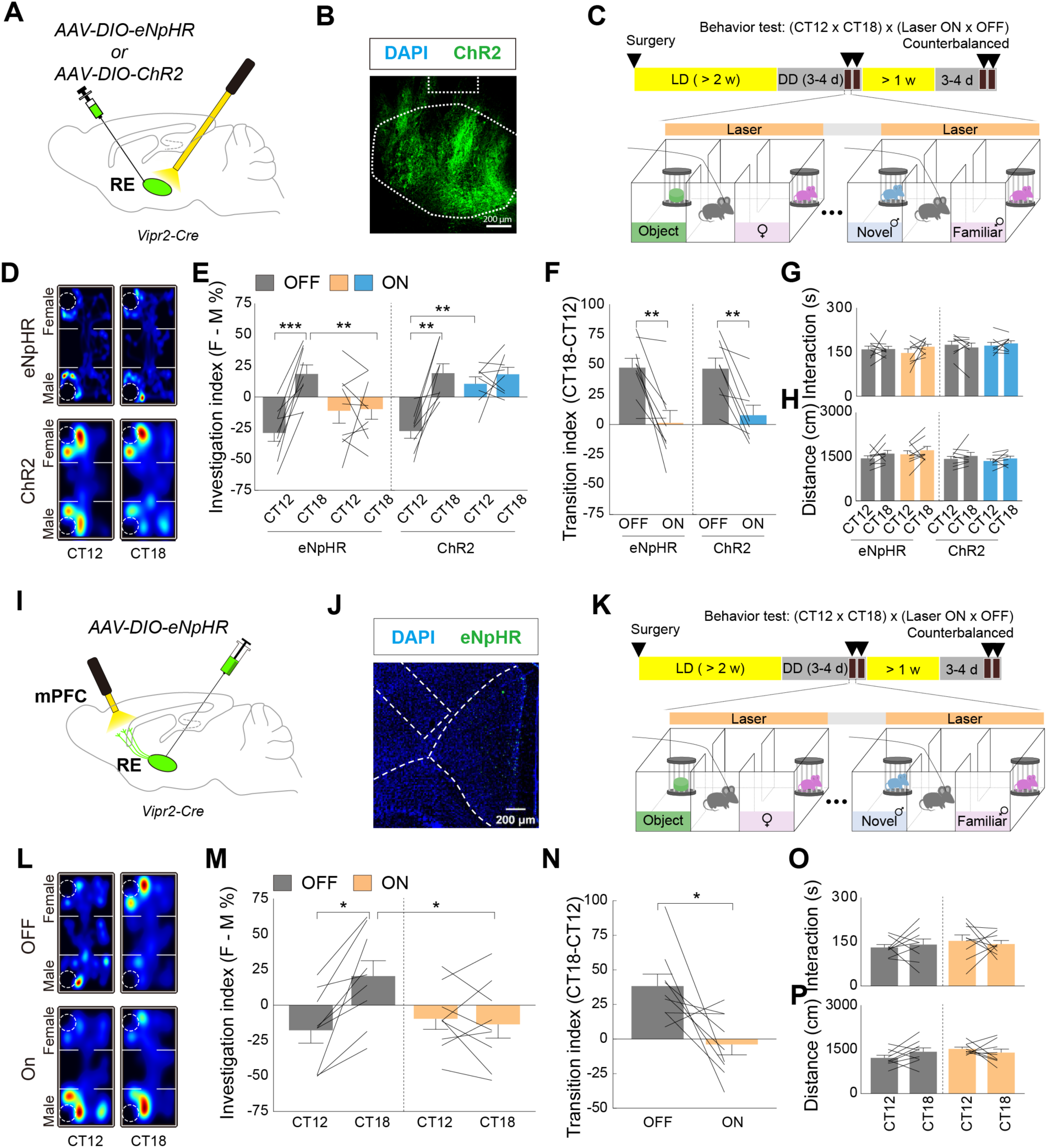
RE^VIPR2^ neuron and its projection to mPFC control circadian rhythm in social priority. (**A** to **B**) Schematics for viral delivery (A) and an example image showing viral expression and optic fiber placement (B). (**C**) Experimental scheme of social priority assay with optogenetic inhibition. (**D**) Representative heatmap visualization of subject mice during social priority assay with optogenetic inhibition in each time point. (**E** to **H**) Quantification of investigation index (E), transition index (F), total interaction (G), and total distance moved (H) during social priority assay with or without optogenetic inhibition. (**I** to **J**) Schematics for viral delivery (I) and an example image showing viral expression and optic fiber placement (J). (**K**) Experimental scheme of social priority assay with optogenetic inhibition. (**L**) Representative heatmap visualization of subject mice during social priority assay with optogenetic inhibition in each time point. (**M** to **P**) Quantification of investigation index (M), transition index (N), total interaction (O), and total distance moved (P) during social priority assay with or without optogenetic inhibition. n = 9 for RE^VIPR2^-eNpHR3.0 animals and n = 8 for RE^VIPR2^-ChR2 animals in (E to H). n = 9 for RE^VIPR2^-mPFC-eNpHR3.0 animals. For (E), (G), (H), (M), (O), and (P), RM Two-way ANOVA test followed by pairwise paired t-test or non-parametric RM Two-way ANOVA test followed by Wilcoxon Signed-Rank test; for (F) and (N), paired t-test was performed. Each line represents a repeated experiment within individual animal. Data are presented as mean ± SEM. \*\*\**P* < 0.001; \*\**P* < 0.01; \**P* < 0.05.

We further explored the downstream target of RE^Vipr2^ neurons by expressing EGFP in RE Vipr2 neurons (fig. S10A, B). RE^Vipr2^ neurons projected predominantly to the mPFC, with minimal projections to the hippocampus. We then optogenetically inhibited this neural circuit during the social priority test (Fig. 4I‒K; fig. S10C, D). Inhibiting RE^Vipr2^-mPFC terminal blocked familiar female investigation at CT18 (Fig. 4L, M) and eliminated the circadian transition in social priority between CT12 and 18 (Fig. 4N), without affecting total interaction time, distance travelled, or behavior during familiarization (Fig. 4O, P; fig. S10E‒G). Collectively, these findings indicate that RE^Vipr2^ neurons and their projections to mPFC shape circadian rhythm in social priority.

### Circadian social priority in naturalistic triadic dynamics

Next, we investigated whether the neural mechanisms described above drive social behavior in a freely interacting triad. We developed another behavioral test, called the three-mice interaction (TMI) assay (Fig. 5B), wherein a subject and two target mice interact freely in an open arena. The TMI paradigm parallels the social priority assay. A subject male mouse was first familiarized with a novel female mouse, after which a second male mouse, novel to both the subject male and the female, was introduced. This configuration enabled simultaneous assessment of interactions between the subject male and the familiar female and between the subject male and the novel male. Social and sexual behaviors of the subject were continuously recorded and analyzed.

**Fig. 5.**
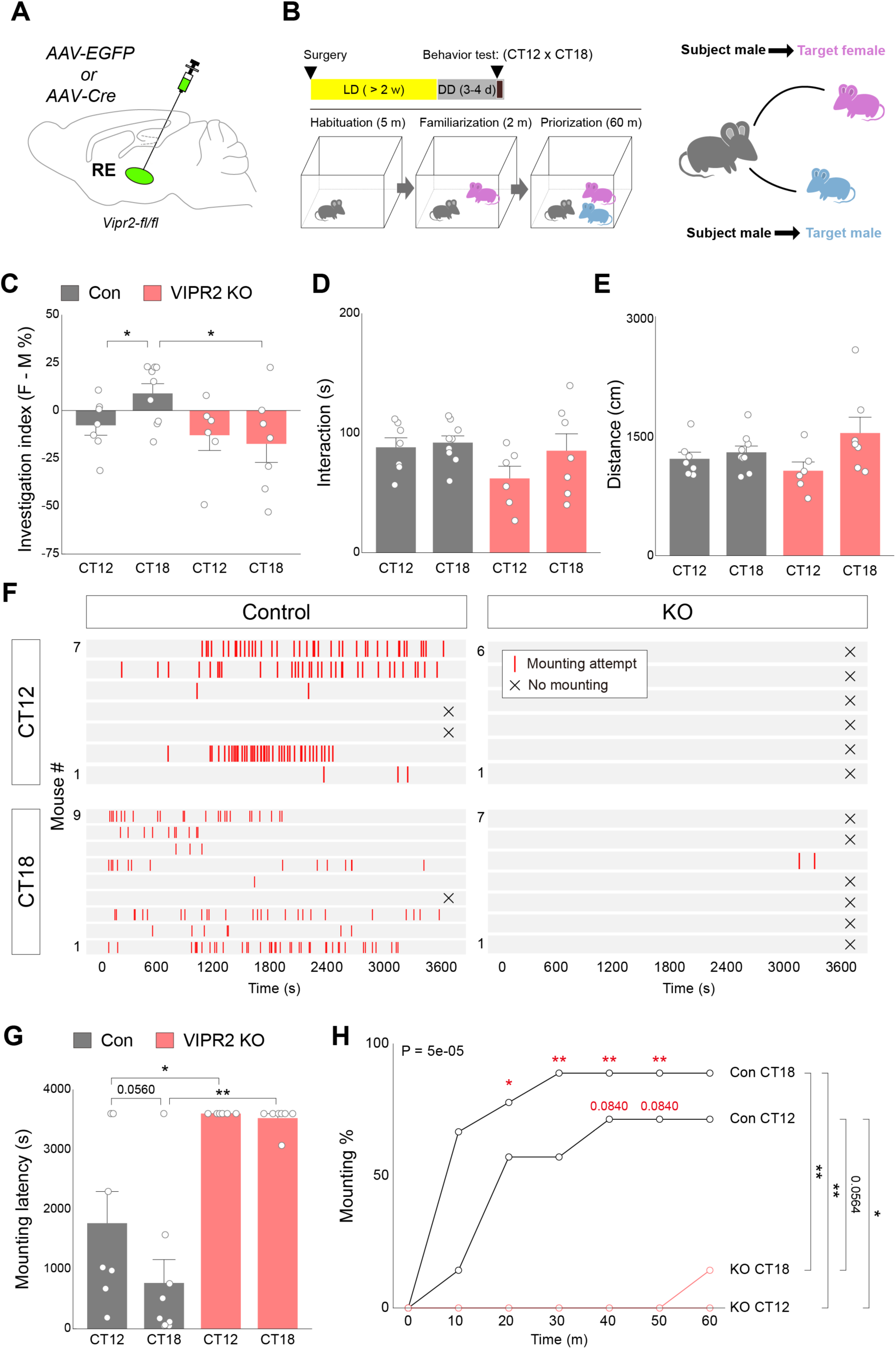
Functional expression of Vipr2 in RE is essential for time-dependent naturalistic group interaction and reproductive behavior. (**A**) Schematics for viral delivery. (**B**) Experimental scheme of TMI assay (left) and illustration of group dynamics in TMI assay (right). (**C** to **E**) Quantification of investigation index (C), total interaction (D), and total distance moved (E) during TMI assay with or without genetic ablation of Vipr2 in RE. (**F**) Rastor plot of subject animals illustrating distribution of mounting attempts during TMI assay. (**G**) Latency to first mounting attempts in each time point with or without genetic ablation of Vipr2 in RE. (**H**) Cumulative plot for the proportion of animals displaying mounting attempts in each time bin. n = 7, 9 for RE-EGFP CT12 and CT18 groups and n = 6, 8 for RE-Cre CT12 and CT18 groups. For (C to E) and (G), Two-way ANOVA test followed by pairwise independent-sample t-test or non-parametric Two-way ANOVA (ART) followed by pairwise Wilcoxon Rank-Sum test; for (H), Log-Rank test followed by pairwise Log-Rank test and Fisher’s exact test followed by pairwise Fisher’s exact test were performed. Each dot represents an individual animal. Data are presented as mean ± SEM. = \*\**P* < 0.01; \**P* < 0.05.

To assess the role of Vipr2, we selectively deleted Vipr2 in the RE by injecting AAV-Cre into the RE of Vipr2fl/fl mice (RE-Vipr2KO) (Fig. 5A). We performed the TMI assay at CT12 and CT18. The appetitive approach toward the familiar female and the novel male quantitatively differed by the time of the day. The investigation index of EGFP-injected control showed enriched investigation toward the novel male (social novelty prioritized) at CT12 and toward the familiar female (sexual preference prioritized) at CT18 (Fig. 5C), consistent with results from the social priority test. Conversely, RE-Vipr2KO consistently favored investigation of the novel male at both time points (Fig. 5C). Neither total interaction time nor distance travelled differed across time points and genotypes (Fig. 5D, E). As an internal control, we analyzed the behavior of target male mice during the TMI assay (fig. S11A). Unlike the subject mice, target male mice were simultaneously exposed to both mice, eliminating social novelty differences. Target male mice showed no circadian changes in social priority, suggesting the interplay between sexual and social novelty preference (fig. S11B‒D). Collectively, these results suggest that the Vipr2 expression in the RE is required for circadian regulation of social priority in a freely behaving triad.

Finally, we explored how circadian rhythm modulation of social priority during the appetitive phase influences consummatory social behavior, specifically reproduction. We continuously recorded direct social interactions among the three mice following the TMI assay. In control animals, mounting attempts by subject males were more frequent at CT18 than at CT12 (Fig. 5F), and mounting latency was significantly shorter at CT18 (Fig. 5G), indicating enhanced sexual behavior when sexual preference is prioritized. However, RE-Vipr2KO mice rarely attempted mounting at both time points (Fig. 5F, G), and instead permitted mounting attempts by the intruder male (fig. S11E, F). Cumulative analyses revealed robust increases in mounting behavior in control mice as well as target male mice, whereas RE-Vipr2KO mice rarely exhibited mounting attempts (Fig. 5H; fig. S11G). Together, the data indicate that Vipr2 signaling in the RE regulates the circadian patterns of both appetitive and consummatory phases of social behaviors.

## Discussion

Social behavior is dynamically shaped by internal states and the identity of interacting partners, yet the neural circuit mechanisms governing these complex decisions remain poorly understood. A major limitation in the field is the predominant use of paradigms centered on single social goals, hindering a comprehensive understanding of how the brain translates simple behavioral control into the richness of real-world social interactions. Here, we address this gap by investigating social prioritization in contexts involving three interacting animals. Our findings reveal a top‒down regulatory mechanism in higher brain centers, as behavioral choices shifted between the early and mid-active phases despite stable capacities for social recognition and sexual approach. This circadian prioritization was further confirmed in a freely moving triad TMI assay, in which preferences for social novelty and sexual interaction were similarly time-of-day dependent. Critically, we identify VIP‒VIPR2 signaling in the RE as the driver of priority transitions across both assays. Therefore, we propose that the interplay between the master clock-thalamocortical pathway and local VIP‒VIPR2 signaling is a key neural principle governing the circadian orchestration of social priorities.

What is the function of circadian social priority between social novelty preference and sexual preference? Circadian rhythmicity is prominent within the reproductive system, where events such as the surge in the hormonal levels of hypothalamus-pituitary-gonad axis and ovulation exhibit distinct circadian patterns (*28*, *29*). These rhythms create temporal windows with superior reproductive efficiency (*17*, *30*). Considering the long refractory period in mice, aligning reproductive behavior with optimal reproductive physiology via circadian regulation is likely advantageous. The circadian regulation of social behavior is particularly pronounced during the consummatory phase, evidenced by the clear circadian patterns of reproductive and aggressive behaviors. The circadian pattern of social priority can therefore provide a temporal filter to match the types of social behavior from appetitive to consummatory phases. Paradoxically, however, sexual preference assays for a single enclosed partner in a three-chamber arena do not show a discernible circadian pattern. The detection of circadian patterns using our social priority test with two enclosed partners highlights the limitations of overly reductionist behavioral models in understanding the mechanisms of social regulation.

Circadian rhythms profoundly impact physiological functions, motivation, emotion, and higher brain functions (*7*, *31*, *32*). However, despite extensive characterization of circadian control over physiology, motivation, and emotion, the mechanisms governing higher-order cognition remain elusive. Our study discovers a functional novel circuit where VIP neurons of the master clock (SCN) communicate with the mPFC through the RE, representing the first demonstration of a neural circuit through which the SCN can exert temporally precise excitatory influence over the prefrontal cortex. We propose that this pathway provides a sophisticated mechanism for fine-tuning cognitive processes. Specifically, the dynamic changes in RE VIPergic tone during the active phase can orchestrate the nuanced, time-of-day-dependent modulation of higher brain functions.

By incorporating circuit-level connections to the master clock (SCN), our study further elucidates the role of the RE in coordinating instinctive social behaviors. These findings highlight the importance of circadian regulation in organizing social instincts across the daily cycle and call for further examinations of how circadian rhythm impacts various instinctive behaviors.

## Acknowledgments

The authors acknowledge Editage (www.editage.com) for English language editing and proofreading. Additionally, Gemini was utilized for initial manuscript refinement.

## Author contributions

Conceptualization: JK, HKC, Methodology: JK, JY, IP, CS, YL, JCR, HKC, Investigation: JK, JY, BK, MK, IP, JL, MC Visualization: JK, JY, HKC, Funding acquisition: HKC, Project administration: JK, HKC, Supervision: KK, JCR, HKC, Writing – original draft: JK, HKC, Writing – review & editing: JK, HKC

## Data and materials availability

All data are available in the main text or the supplementary materials.

## Supplementary Materials

Materials and Methods

Supplementary Text

Figs. S1 to S11

Tables S1

References (*1–38*)

Data S1

