## Supplementary materials for "Master clock‒thalamic‒prefrontal circuit controls circadian social priority"

Science
AAAS

### Supplementary Materials for

#### Master clock-thalamic-prefrontal circuit controls circadian social priority

Jihoon Kim[1,2], Jinseon Yu[1,4], Boil Kim[1], Inah Park[1,2], Minseok Kim[1,3], Jimin Lee[1], Mijung Choi[1,2], Cheol Song[5], Yulong Li[6], Kyungjin Kim[1], Jong-Cheol Rah[1,4], Han Kyoung Choe[1,2,3]*

**The PDF file includes:**

Materials and Methods
Figs. S1 to S11

#### Materials and Methods

Animals

All procedures were approved by the Institutional Animal Care and Use Committee of the Daegu Gyeongbuk Institute of Science and Technology (DGIST). Adult male C57BL6/J, *Vip-ires-Cre* (#010908, Jackson Laboratory), *Avp-ires-Cre* (#023530, Jackson Laboratory), *Vipr2-floxed* (#035395, Jackson Laboratory), *Vipr2-ires-Cre* (#031332, Jackson Laboratory), and Ai14 (#007914, Jackson Laboratory) mice were reared in standard cages (16 × 36 × 12.5 cm3) and housed under 12-h light/dark cycle (light on from 01:00 to 13:00) with water and food available ad libitum. Mice were weaned from same-sex littermates at approximately 3–4 weeks of age. Mice aged 8–10 weeks were used in the experiments. All behavioral experiments were performed under dimly lit red light conditions.

Adeno-associated virus (AAV) production

AAV vector expressing $GRAB_{VIP1.0}$ was packaged as previously described, with modifications (*33*). After AAV production, their genomic copies were quantified using real-time quantitative polymerase chain reaction. The AAVs were diluted in phosphate-buffered saline (PBS) to a final concentration of 8.12 X $10^{12}$ genomic copies per milliliter (GC/mL) before the experiments.

AAV and stereotaxic surgery

AAVs used for the experiments are as follows: AAV1-Ef1a-DIO-eNpHR3.0-EYFP (GC/mL; Addgene), AAV1-Ef1a-DIO-ChR2-EYFP (GC/mL; Addgene), AAV1-Ef1a-DIO-EGFP (GC/mL; Addgene), AAV2-hSyn-Cre::EGFP (GC/mL; Addgene), AAV2-hSyn-EGFP (GC/mL; Addgene), AAV1-Ef1a-DIO-GCaMP7s (GC/mL; Addgene), AAV1-CAG-DIO-$GRAB_{VIP1.0}$ (8.12 X $10^{12}$ GC/mL).

The mice (8–10 weeks old) selected for the experiments were anesthetized with a mixture of ketamine (100 mg/kg) and xylazine (10 mg/kg) in 0.9% saline and placed in a stereotaxic frame (RWD Life Science) for viral injection. After disinfection and craniotomy above the target region, the virus was injected using a nanoliter injector system (World Precision Instruments) at an injection rate of <100 nL/min. The coordinate for each brain regions are as follows: For SCN, AP: -1.0 mm, ML: ±0.25 mm, DV: -5.75 mm; For 20° angled SCN injection, AP: -1.0 mm, ML: ±2.25 mm, DV: -5.5 mm; For the nucleus reuniens (RE), AP: -1.35 mm, ML: ±0.25 mm, DV: -4.6 mm; For 20° angled RE injection, AP: -1.0 mm, ML: ±1.62 mm, DV: -4.65 mm; For 20° angled PVN injection, AP: -1.3, ML: ±1.96 mm, DV: -4.75 mm; For 20° angled MPOA injection, AP: -0.45 mm, ML: ±2.09 mm, DV: -4.55 mm; For PVT, AP: -0.9 mm, ML: ±0.25 mm, DV: -3.3 mm; For lateral ventricle (LV), AP: -0.5 mm, ML: -1.0 mm, DV: -1.4 mm; For 20° angled mPFC injection, AP: +1.7 mm, ML: ±1.3 mm, DV: -1.0 mm. Optic fibers (400 µm for single fiber implantation and 200 µm for bilateral, angled fiber implantation; 0.5 NA; Neurophotometrics) were placed ~100 µm above viral injection coordinates. Cannula (outer diameter, 0.48 mm; inner diameter, 0.34 mm; #62003, RWD Life Science) was implanted in the LV for pharmacological experiments.

Intracerebroventricular (I.C.V.) injection

VIP receptor (VIPR) antagonist ([D-p-Cl-$Phe^6$, $Leu^{17}$]-VIP; #3054, Tocris), VIPR1 selective antagonist (PG 97-269; Bachem), and VIPR2 selective antagonist (PG 99-465; Bachem) were prepared in artificial cerebrospinal fluid (aCSF; #35-252, Tocris) in 1 mM concentration. 2 μL of antagonists were infused into the LV at a speed of 1 μL/min. After I.C.V. injection, mice were returned to the home cage for 30 min before the experiment.

Vaginal cytology for the estrus cycle

Before initiating the experiment, the estrus cycle of each female mouse was confirmed via vaginal cytology. Briefly, 200–300 μL of 0.9% saline was aspirated with a disposable pipette, and the vaginal area was rinsed approximately 10 times. Subsequently, 0.9% saline solution containing cells from the vaginal epithelium was examined under the microscope, and the estrus phase of each animal was confirmed via visual inspection of the morphology of epithelial cells and neutrophils. Samples containing neutrophils (metestrus or diestrus) were excluded from the experiment, and female mice with nucleated or anuclear epithelial cells (proestrus or estrus) were used for further experiment.

Behavioral assays

All behavior assays were performed at indicated time of day under dimly lit red light conditions, following three days of constant dark (DD) condition as previously described (*34*).

*Three-chamber assay*

Three-chamber assay was performed as previously reported with some modifications (*35*). Three-chamber apparatus was prepared with doors closed and empty wire cups placed on either side. The subject mouse was placed in the center chamber of the three-chamber apparatus and allowed habituation to the experimental setting for 5 min (center habituation). After center habituation, side doors were removed for free access to either chamber for 5 min (three-chamber habituation). The three-chamber apparatus was cleaned with 70% ethanol between each session. We further conducted each three-chamber-based tests as follows.

For sociability test, after three-chamber habituation, doors were placed again with the subject mouse in the center chamber, and social and non-social cues (a male mouse and a mouse-sized object) were placed inside the wire cups on either side. After a brief interval (2 min), doors were removed, and the free exploration of the subject mouse between the two side chambers was measured. The three-chamber apparatus was cleaned with 70% ethanol between each session.

For social recognition test, after completing sociability measurements, doors were placed again with the subject mouse in the center chamber, and the non-social cue was removed, and another social cue (a male mouse) was placed inside the wire cup. After a brief interval (2 min), doors were removed, and the free exploration of the subject mouse between familiar and novel social cues was measured. For the social novelty preference test for the female subject mice, two female mice were used as the target cue.

For sexual preference test, after completing center habituation and three-chamber habituation, male and female social cues (a male and a female mouse) were placed inside the wire cups on either side. After a brief interval (2 min), doors were removed, and the free exploration of the subject mouse between the two side chambers was measured.

For social priority test, after completing center and three-chamber habituation, social and non-social cues (a female mouse and a mouse-sized object) were placed inside the wire cups on either side. After a brief interval (2 min), doors were removed, and the subject mouse was allowed to freely explore between the two chambers (familiarization). After completing the familiarization phase, doors were placed again with the subject mouse in the center chamber, the non-social cue was removed, and another social cue (a male mouse) was placed inside the wire cup. After a brief interval (2 min), doors were removed, and the free exploration of the subject mouse between novel male cue and familiar female cue was measured (prioritization). For the social prioritization test for the female subjective mice, male cue was used during familiarization, and familiar male and novel female cues were used during prioritization.

*Three-mouse interaction (TMI) assay*

TMI assay was conducted in a square arena with sufficient bedding. Target male and female mice were labeled with a marker pen on their tail to distinguish each animal during subsequent video analysis. The subject mouse was placed inside the arena and allowed free exploration for 5 min (habituation). After, a female mouse was introduced into the arena for 2 min for direct interaction between the subject and the target mouse. After the end of the 2-min interaction, another male mouse was introduced into the arena, and the free interaction between the three mice was recorded for 60 min. The interaction time between each animal was measured for the first 5 min of the three-mouse recording, and reproductive behavior of both the subject and the target mouse was measured for 60 min.

*Behavior analysis*

Each behavior was recorded via a camera connected to EthoVision (Noldus) software. For the three-chamber assays, the duration of each animal around the wire cups (sniffing zone) was measured, and the investigation index was calculated as: ($sniffing\ zone\ duration^{target\ 1} - sniffing\ zone\ duration^{target\ 2}$) / ($sniffing\ zone\ duration^{target\ 1} + sniffing\ zone\ duration^{target\ 2}$) X 100. For quantification of the investigatory behavior during the TMI assay, 5-min recordings of each group were manually examined and corrected using EthoVision software. The interaction time was calculated by thresholding the distance between the body parts of each animal within 6 cm as “interaction.” For quantification of reproductive behavior during the TMI assay, the mounting attempt (trying to climb over the back of the female mouse with hind legs repeatedly kicking off the ground) was manually measured, and a mounting bout with thrusting behavior was labeled as mounting success. For the experiment with repeated measures between circadian time points, the transition index was calculated as the subtraction of the investigation index between those time points.

*Optogenetic manipulation*

Mice were acclimated to the handling, patch cord attachment, and three-chamber exploration (30 min) with the patch cord attachment 1–2 days before the experiment. For optogenetic inhibition, a continuous laser with a 589-nm wavelength (MSL-FN-589; CNI laser) was applied with 10–20 mW intensity at the tip of the optic cannula. The same laser intensity was kept for both 200- and 400-µm optic cannula. For optogenetic activation, a laser with a 470-nm wavelength (MBL-FN-473; CNI laser) was applied at 10 mW intensity and 30 Hz frequency with 15 ms pulse width. Both optogenetic inhibition and activation were applied only during familiarization and prioritization sessions of social prioritization.

<u>Fiber photometry</u>

*Calcium activity recording*

Mice were acclimated to the handling, patch cord attachment, and three-chamber exploration (30 min) with the patch cord attachment 1–2 days before the experiment. The intensity of the 465-nm LED was kept around 20–30 µW at the tip of the optic cannula. Recording was conducted during the social prioritization assay, with the start of each session in the assay time-stamped using a transistor–transistor logic output signal from the EthoVision software.

*VIPR activity recording*

Mice were acclimated to the handling and patch cord attachment 1–2 days before the experiment. On the day of recording, the subject mouse was attached to the path cord with a 465-nm LED at 10–20 µW and placed into a square arena with free access to food and water and sufficient bedding. The fluorescence was measured continuously throughout 24 h.

*Data analysis*

The fiber photometry data were processed as previously described (*36*). Briefly, 465 (F465) and 405 (F405) nm LEDs were used for calcium-dependent and -independent GCaMP7s fluorescence excitation, and the emitted signal was passed through a low-pass filter and demodulated using Synapse software (Tucker-Davis Technologies). The F405 signal was used to correct motion artifacts and was normalized using the formula dF/F = (F465 – fitted F405)/fitted F405. To compare calcium response between experimental animals, global Fz was calculated as Fz = (dF/F – median(dF/F))/std(dF/F).

For peri-event histogram analysis for each social cue, the dF/F signal was sliced and aligned to event onsets. The signal before the event onset was used as the baseline (FB) for z-normalization. Fz was calculated as Fz = (dF/F – mean(FB))/std(FB). To quantify the global calcium level, the mean and standard deviation of the entire session were used: Fz = (dF/F – mean(dF/F))/std(dF/F).

For VIPR activity recording, fluorescence decay was corrected and smoothed using 'msbackadj' before calculating dF/F. dF/F was median-filtered and CT11 to CT12 window was used as the baseline for z-normalization.

<u>Electrophysiological recording</u>

*Slice preparation*

Acute slices were prepared from VIPR2-IRES Cre mice. Transcardial perfusion was performed after anesthetizing the mice with sodium pentobarbital at 70 mg/kg. The brains were rapidly removed and immersed in chilled cutting solution of the following composition (in mM): 110 choline chloride, 2.5 KCl, 25 $NaHCO_3$, 1.25 $NaH_2PO_4$, 25 glucose, 0.5 $CaCl_2$, 7 $MgCl_2{\cdot}6H_2O$, 11.6 sodium L-ascorbate, and 3 pyruvic acid. Thereafter, 300 µm-thick coronal slices were prepared using a vibratome (VT1200S, Leica, Wetzlar, Germany). The slices were then incubated for 30 min at 32°C in aCSF containing the following (in mM): 119 NaCl, 2.5 KCl, 26 $NaHCO_3$, 1.25 $NaH_2PO_4$, 20 glucose, 2 $CaCl_2$, 1 $MgSO_4$, 0.4 L-Ascorbic acid, and 2 pyruvic acid. During sectioning, the solutions were saturated with carbogen (95% $O_2$, 5% $CO_2$) to a final pH of 7.4.

*Electrophysiological recording*

The slices were transferred to a submerged recording chamber with a continuous flow of aCSF saturated with carbogen (95% O2, 5% CO2) for whole-cell patch-clamp recordings. RE $VIPR2^+$ neurons were visualized using an upright microscope (BX51WI, Olympus) equipped with differential interference contrast optics with a 40x water immersion objective (NA 0.8, Olympus). All electrophysiological recordings were performed at 30 ± 2°C, and fresh aCSF was perfused at approximately 1.5 mL/min. The patch electrodes were pulled from borosilicate glass capillaries to obtain a resistance between 4 and 5 MΩ. The internal solution contained the following (in mM): 138 potassium gluconate, 10 KCl, 10 HEPES, 10 $Na_2$-phosphocreatine, 4 MgATP, 0.3 NaGTP, and 0.2 EGTA (pH 7.25).

*Drug application*

VIP (1 µM; Tocris Bioscience, Bristol, UK; Cat. No. 1911), mixed with Alexa 594 for visualization, was dissolved in aCSF and loaded into a glass pipette (tip diameter: 1–2 µm). To avoid mechanical displacement of the tissue by pressure, the pipette tip was positioned slightly away from the recorded cell. Puff application was delivered using pressure pulses (30 kPa) lasting from 100 to 5,000 ms.

Confocal imaging and analysis

Mice were perfused and fixed in 1X PBS and 4% paraformaldehyde (PFA) after blood collection. Brains were isolated and post-fixed in 4% PFA at 4°C overnight. Following post-fixation for 24 h, brains were either incubated in 30% sucrose and then frozen in an OCT compound or in PBS. Brains were sectioned at a thickness of 50 µm using a cryostat or vibratome, and every sixth section was used for subsequent analysis. The sections were stained with DAPI (1:1,000) before mounting on a glass cover. Images were obtained using a Nikon C2+ confocal microscope.

Statistics

Statistical analyses were performed using R software (version 4.2.3). Levene's test and Shapiro-Wilk test were performed before further statistical analysis and used as the basis for the appropriate method. A paired t-test was used for repeated experiments with two groups. An unpaired t-test was used for independent experiments with two groups. One-way analysis of variance (ANOVA) or Kruskal–Wallis test followed by post hoc Tukey's Honestly Significant Difference test was used to compare multiple groups. Two-way ANOVA or aligned rank transform (ART) ANOVA followed by pairwise independent-sample t-test (Holm-corrected) or pairwise Wilcoxon Rank-Sum test (Holm-corrected) were used for 2X2 factorial design experiments (*37*). Repeated measure Two-way ANOVA followed by post-hoc pairwise paired t-test (Bonferroni-corrected) or nonparametric Two-way ANOVA (nparLD) followed by post-hoc Wilcoxon Signed-Rank test were used for repeated 2X2 factorial design experiments (*38*). Two-way mixed ANOVA followed by post hoc pairwise paired t-test and pairwise independent-sample t-test, or nonparametric Two-way mixed ANOVA (nparLD) followed by post hoc Wilcoxon Signed-Rank test and Wilcoxon Rank-Sum test were used for 2X2 mixed-design experiments. Log-Rank test followed by pairwise Log-Rank test (Holm-corrected) were used to compare the rates of behavior between groups over time. Fisher's exact test followed by pairwise Fisher's exact test (Holm-corrected) were used to compare the distribution of behavioral responses between groups in each time bin.

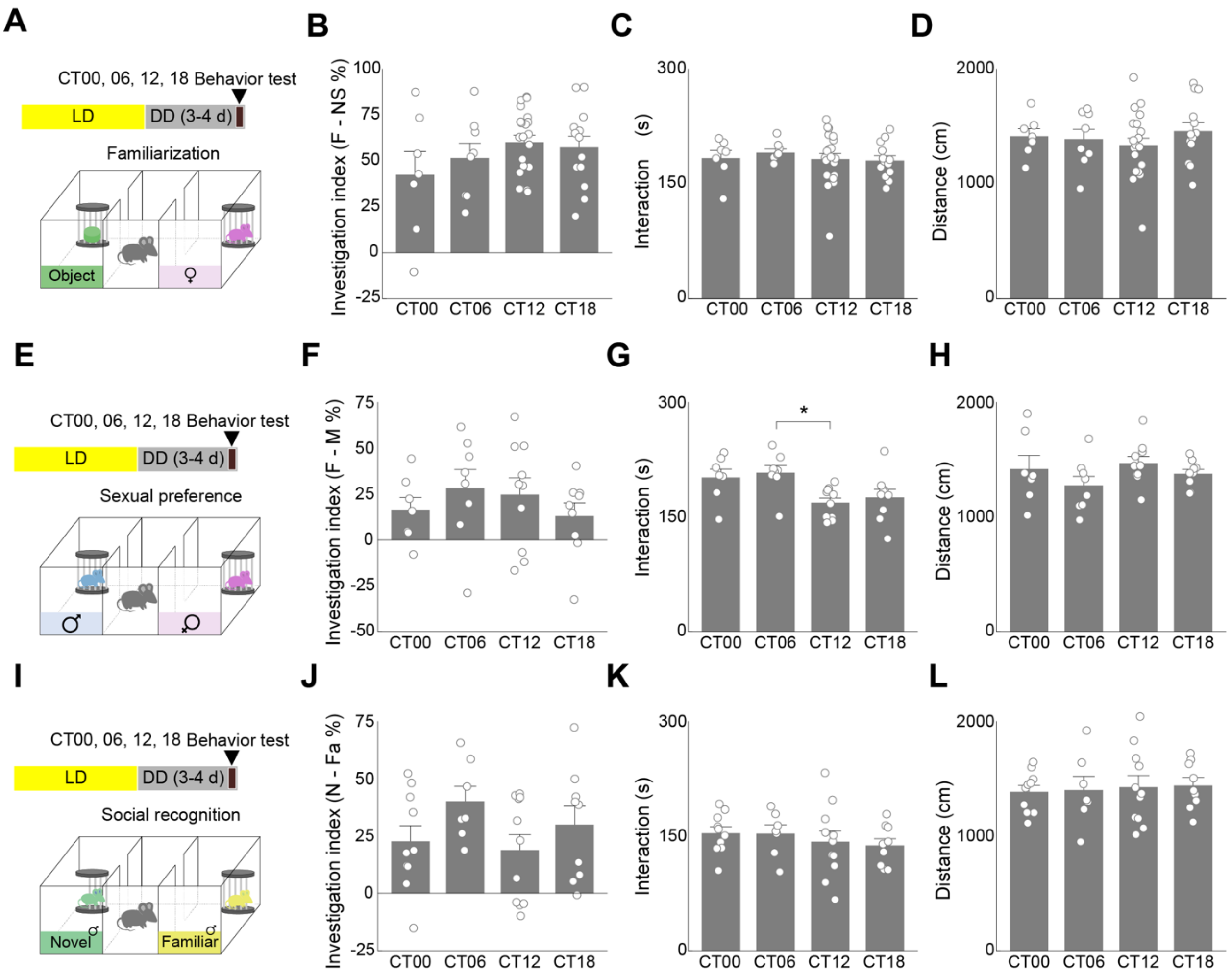

A
B
C
D
CT00, 06, 12, 18 Behavior test
LD
DD (3-4 d)
Familiarization
Object
♀
Investigation index (F - NS %)
Interaction (s)
Distance (cm)
CT00 CT06 CT12 CT18
E
F
G
H
Sexual preference
♂
Investigation index (F - M %)
*
I
J
K
L
Social recognition
Novel ♂
Familiar ♂
Investigation index (N - Fa %)

**Fig. S1. Circadian profile of familiarization, sexual preference, and social recognition.**

(**A**) Experimental scheme of familiarization session of social priority assay. (**B** to **D**) Quantification of investigation index (B), total interaction (C), and total distance moved (D) during familiarization session of social priority assay in each time point. (**E**) Experimental scheme of sexual preference assay. (**F** to **H**) Quantification of investigation index (F), total interaction (G), and total distance moved (H) during sexual preference assay in each time point. (**I**) Experimental scheme of social recognition assay. (**J** to **L**) Quantification of investigation index (J), total interaction (K), and total distance moved (L) during social recognition assay in each time point. n = 7, 8, 21, 13 for CT00, 06, 12, 18 groups in (B to D); n = 7, 8, 10, 9 for CT00, 06, 12, 18 groups in (F to H); n = 10, 7, 11, 9 for CT00, 06, 12, 18 groups in (J to L). For (B to D), (F to H), and (J to L), One-way ANOVA test followed by Tukey's HSD test or Kruskal-Wallis test was performed. Each dot represents an individual animal. Data are presented as mean ± SEM. *$P < 0.05$.

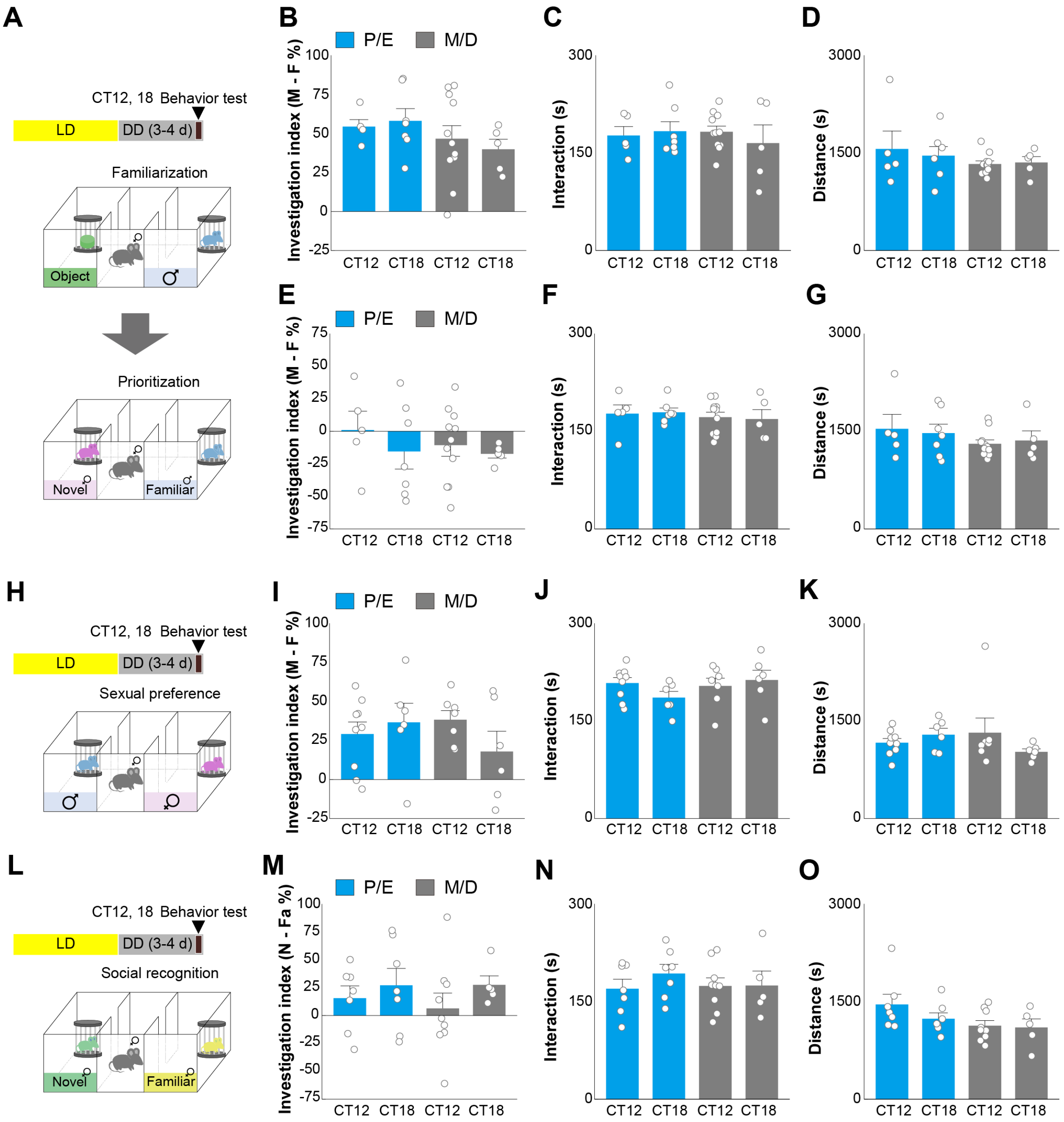
A
CT12, 18 Behavior test
LD
DD (3-4 d)
Familiarization
Object
Prioritization
Novel
Familiar
B
P/E
M/D
Investigation index (M - F %)
CT12 CT18 CT12 CT18
C
Interaction (s)
D
Distance (s)
E
F
G
H
Sexual preference
I
J
K
L
Social recognition
M
Investigation index (N - Fa %)
N
O

**Fig. S2. Circadian profile of social priority, sexual preference, and social recognition in female mice.**

(**A**) Experimental scheme of familiarization and prioritization session of social priority assay. (**B** to **D**) Quantification of investigation index (B), total interaction (C), and total distance moved (D) during familiarization session of social priority assay in each time point. (**E** to **G**) Quantification of investigation index (E), total interaction (F), and total distance moved (G) during prioritization session of social priority assay in each time point. (**H**) Experimental scheme of sexual preference assay. (**I** to **K**) Quantification of investigation index (I), total interaction (J), and total distance moved (K) during sexual preference assay in each time point. (**L**) Experimental scheme of social recognition assay. (**M** to **O**) Quantification of investigation index (M), total interaction (N), and total distance moved (O) during social recognition assay in each time point. n = 5 and 7 for P/E animals, and n = 11 and 5 for M/D animals for CT12 and 18 groups in (B to G); n = 9 and 6 for P/E animals, and n = 7 and 6 for M/D animals for CT12 and 18 groups in (I to K); n = 7 and 7 for P/E animals, and n = 9 and 5 for M/D animals for CT12 and 18 groups in (M to O). For (B to G), (I to K), and (M to O), One-way ANOVA test or Kruskal-Wallis test was performed. Each dot represents an individual animal. Data are presented as mean ± SEM.

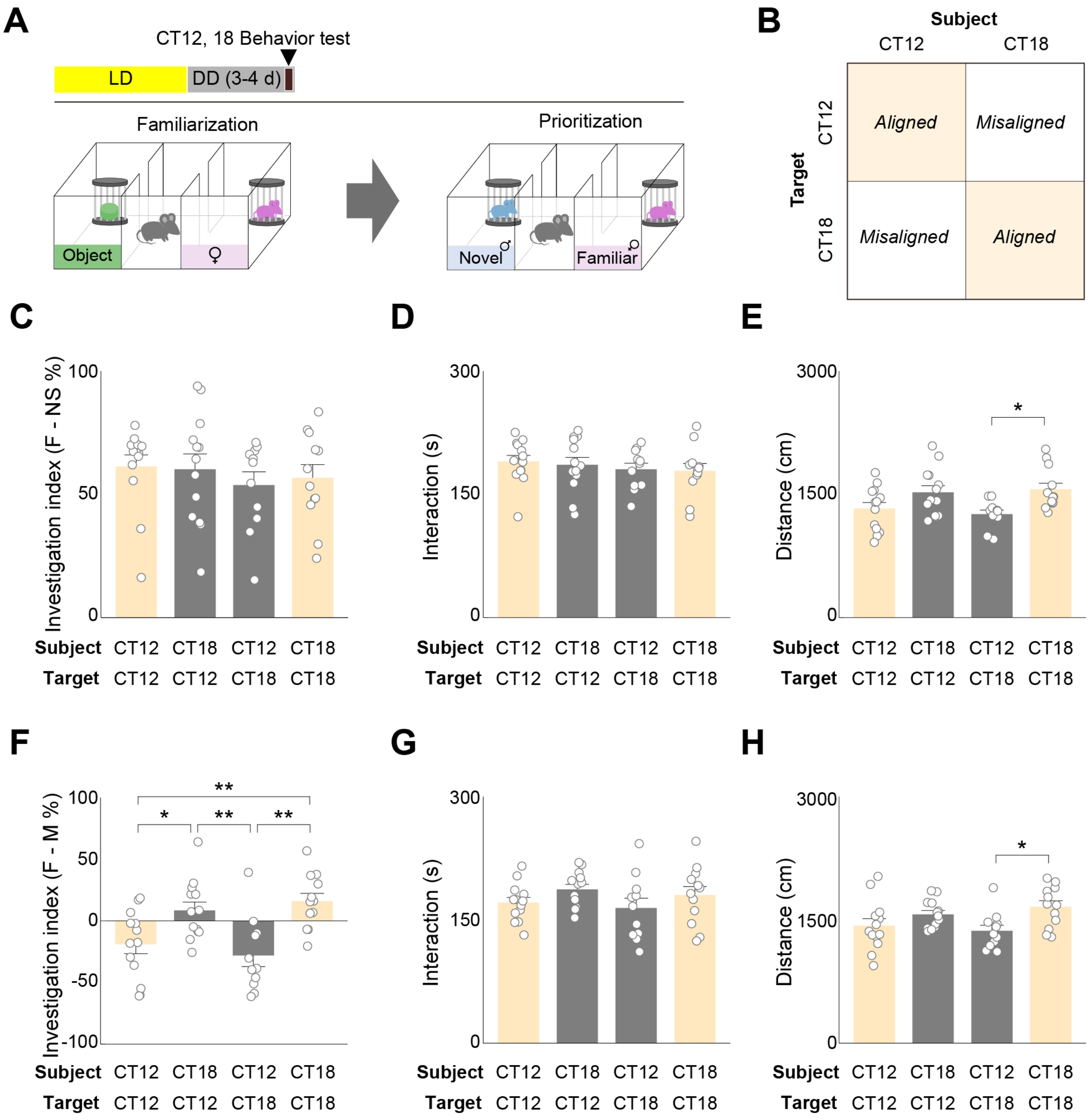
A
CT12, 18 Behavior test
LD
DD (3-4 d)
Familiarization
Prioritization
Object
♀
Novel ♂
Familiar ♀
B
Subject
CT12
CT18
Target
Aligned
Misaligned
Misaligned
Aligned
C
D
E
F
G
H
Investigation index (F - NS %)
Interaction (s)
Distance (cm)
Investigation index (F - M %)
Subject CT12 CT18 CT12 CT18
Target CT12 CT12 CT18 CT18

**Fig. S3. Influence of circadian misalignment on social priority.**

(**A**) Experimental scheme of familiarization and prioritization session of social priority assay. (**B**) Illustration of circadian misalignment between subject and target animals. (**C** to **E**) Quantification of investigation index (C), total interaction (D), and total distance moved (E) during familiarization session of social priority assay in each time point. (**F** to **H**) Quantification of investigation index (F), total interaction (G), and total distance moved (H) during prioritization session of social priority assay in each time point. n = 13 for CT12 aligned, n = 13 for CT18 misaligned, n = 11 for CT12 misaligned, and n = 12 for CT18 aligned animals in (C to H). For (C to H), One-way ANOVA test followed by Tukey's HSD test or Kruskal-Wallis test was performed. Each dot represents an individual animal. Data are presented as mean ± SEM. **$P < 0.01$; *$P < 0.05$.

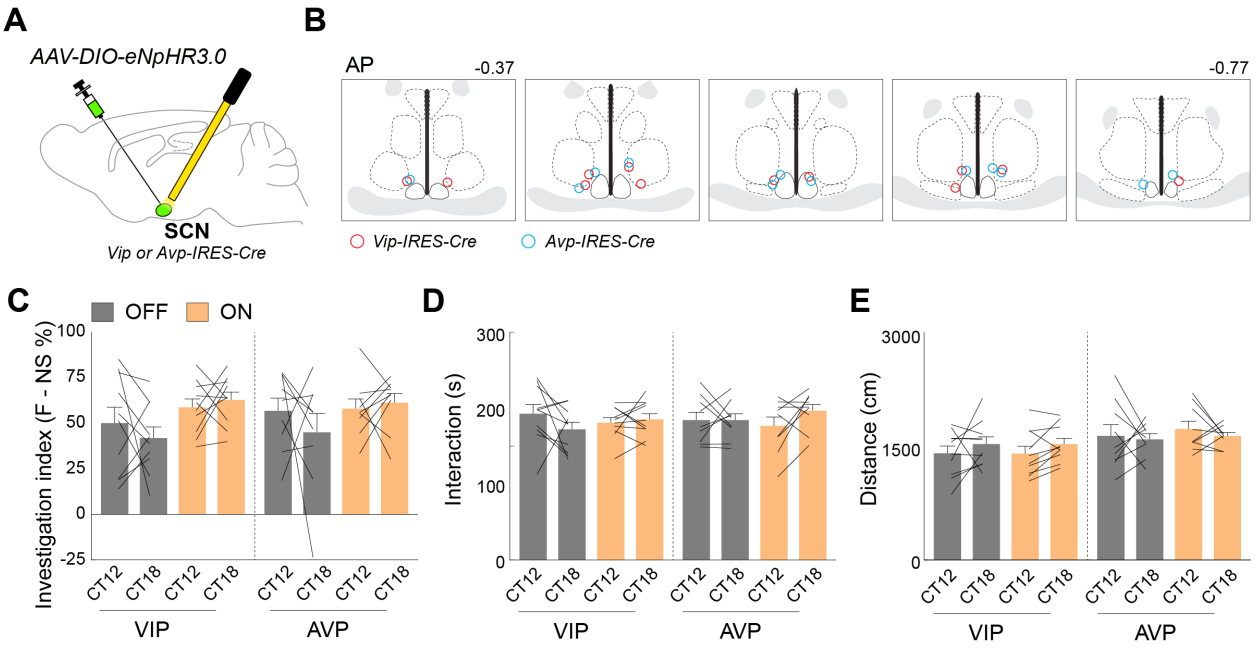

A
AAV-DIO-eNpHR3.0
SCN
Vip or Avp-IRES-Cre
B
AP
-0.37
-0.77
Vip-IRES-Cre
Avp-IRES-Cre
C
OFF
ON
Investigation index (F - NS %)
VIP
AVP
D
Interaction (s)
E
Distance (cm)
CT12
CT18

**Fig. S4. Influence of optogenetic inhibition of SCN neuron during familiarization session.** (**A**) Schematics for viral delivery. (**B**) Illustration of optic fiber placement. (**C** to **E**) Quantification of investigation index (C), total interaction (D), and total distance moved (E) during familiarization session of social priority assay with optogenetic inhibition in each time point. n = 10 for $SCN^{VIP}$-eNpHR3.0 animals and n = 9 for $SCN^{AVP}$-eNpHR3.0 animals in (C to E). For (C to E), RM Two-way ANOVA test or non-parametric RM Two-way ANOVA test was performed. Each line represents a repeated experiment within individual animal. Data are presented as mean ± SEM.

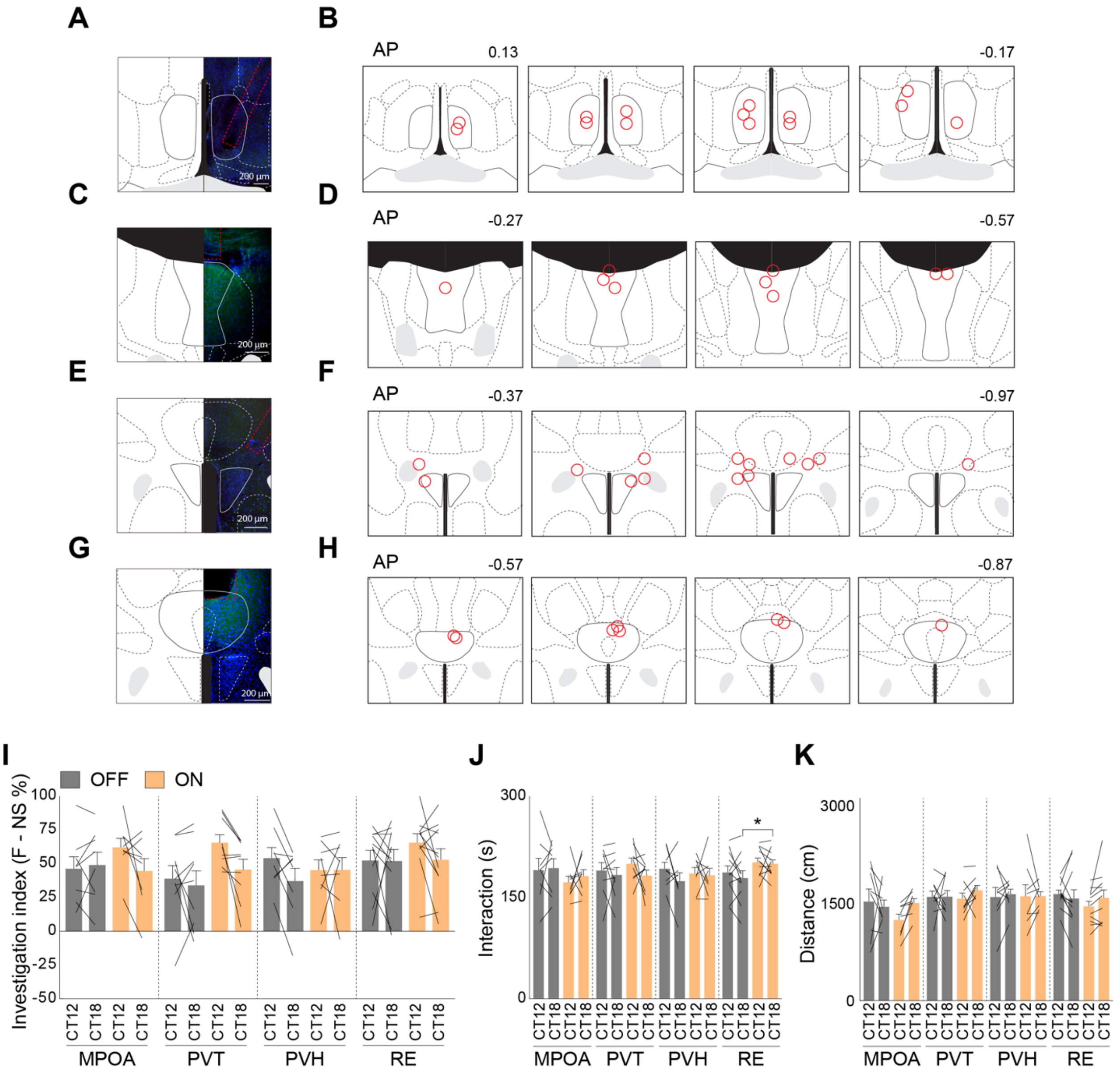

A
B
AP
0.13
-0.17
200 µm
C
D
AP
-0.27
-0.57
E
F
AP
-0.37
-0.97
G
H
AP
-0.57
-0.87
I
OFF
ON
Investigation index (F - NS %)
100
75
50
25
0
-25
-50
CT12
CT18
MPOA
PVT
PVH
RE
J
Interaction (s)
300
150
0
*
K
Distance (cm)
3000
1500
0

**Fig. S5. Influence of optogenetic inhibition of SCN$^{VIP}$ neural terminal during familiarization session.**

(**A**) An example image showing viral expression and optic fiber placement in MPOA region. (**B**) Illustration of optic fiber placement in MPOA region. (**C**) An example image showing viral expression and optic fiber placement in PVT region. (**D**) Illustration of optic fiber placement in PVT region. (**E**) An example image showing viral expression and optic fiber placement in PVN region. (**F**) Illustration of optic fiber placement in PVN region. (**G**) An example image showing viral expression and optic fiber placement in RE region. (**G**) Illustration of optic fiber placement in RE region. (**I** to **K**) Quantification of investigation index (I), total interaction (J), and total distance moved (K) during familiarization session of social priority assay with optogenetic inhibition in each time point. n = 8 for SCN$^{VIP}$-MPOA-eNpHR3.0, n = 9 for SCN$^{VIP}$-PVT-eNpHR3.0, n = 8 for SCN$^{VIP}$-PVH-eNpHR3.0, and n = 11 for SCN$^{VIP}$-RE-eNpHR3.0 animals (I to K). For (I to K), RM Two-way ANOVA test followed by pairwise paired t-test or non-parametric RM Two-way ANOVA test was performed. Each line represents a repeated experiment within individual animal. Data are presented as mean ± SEM.

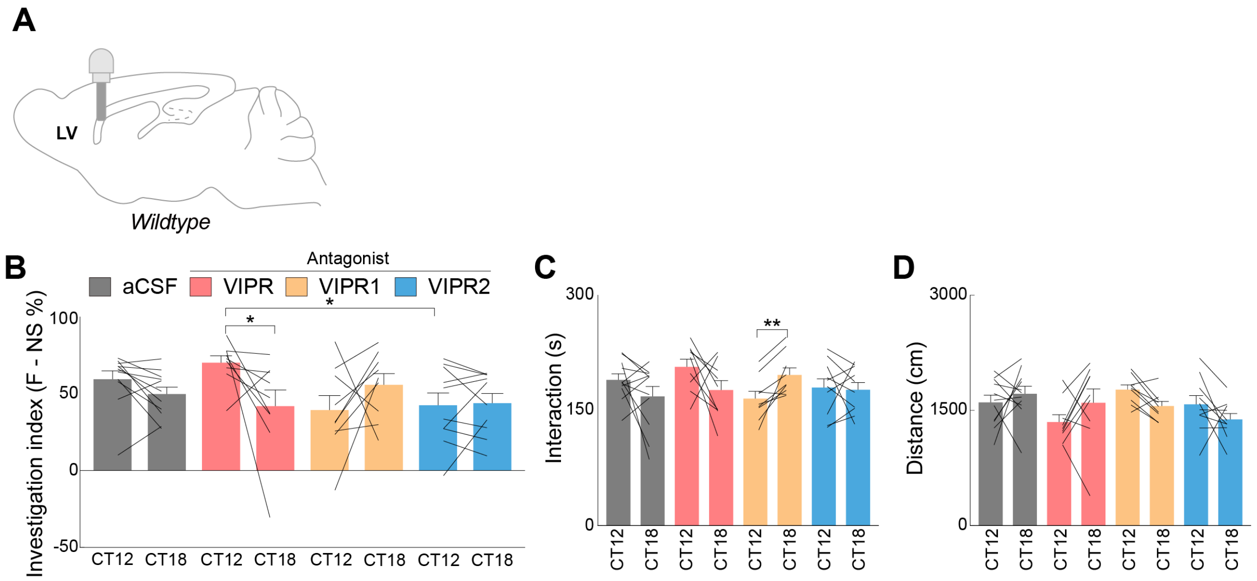
A
LV
Wildtype
B
Antagonist
aCSF
VIPR
VIPR1
VIPR2
Investigation index (F - NS %)
100
50
0
-50
*
*
CT12
CT18
C
300
150
0
Interaction (s)
**
D
3000
1500
0
Distance (cm)

**Fig. S6. Influence of pharmacological inhibition of systematic VIPR signaling during familiarization session.**

(**A**) Schematics for cannula implantation. (**B** to **D**) Quantification of investigation index (B), total interaction (C), and total distance moved (D) during familiarization session of social priority assay with pharmacological inhibition. n = 11, 9, 9, 10 for aCSF, VIPR, VIPR1, and VIPR2 antagonist treatment groups in (B to D). For (B to D), Two-way mixed ANOVA test followed by pairwise paired t-test and pairwise independent-sample t-test or non-parametric Two-way mixed ANOVA test followed by pairwise Wilcoxon Signed-Rank test and pairwise Wilcoxon Rank-Sum test was performed. Each line represents a repeated experiment within individual animal. Data are presented as mean ± SEM. *$P < 0.05$.

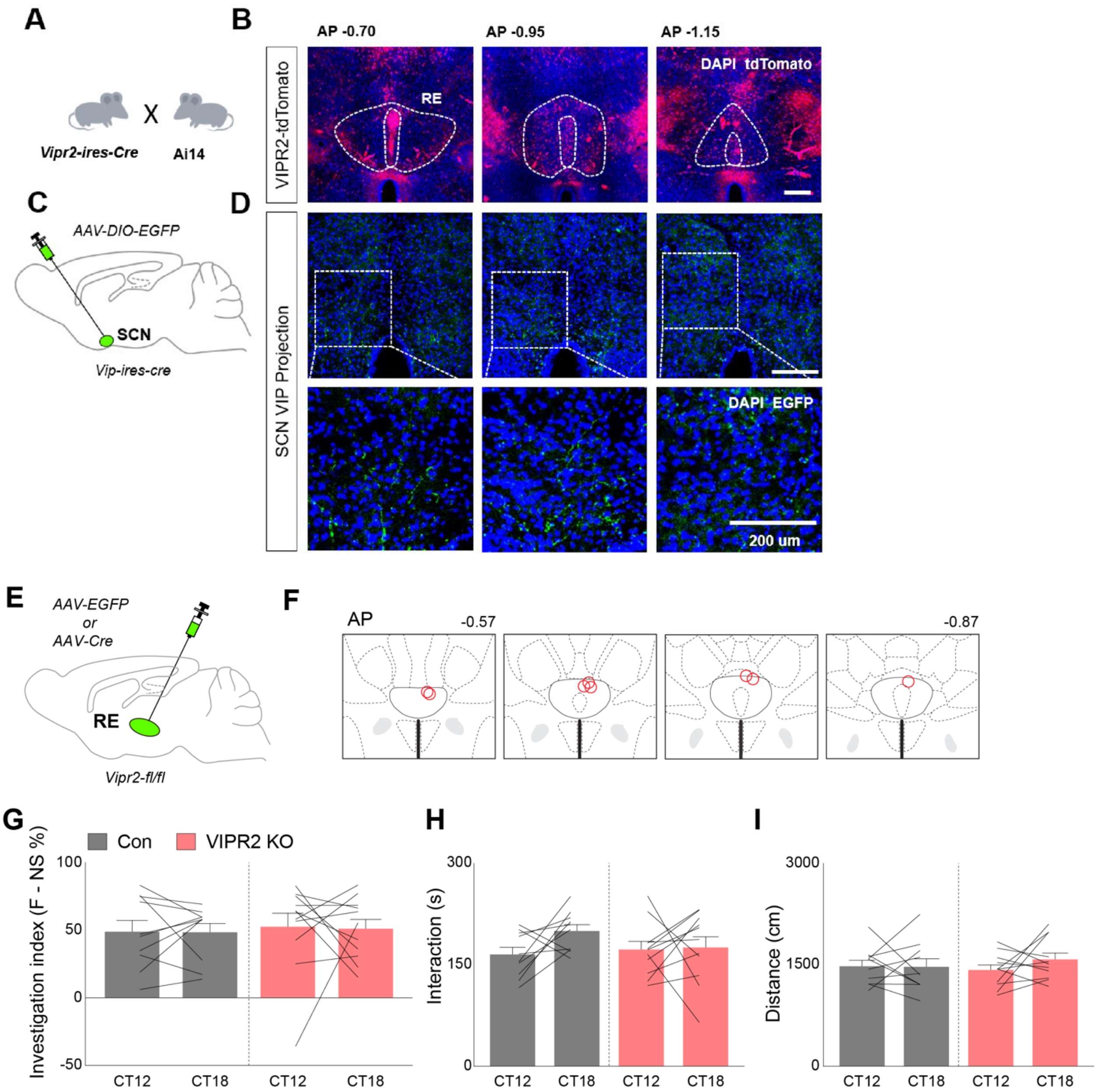
A
Vipr2-ires-Cre
X
Ai14
B
AP -0.70
AP -0.95
AP -1.15
VIPR2-tdTomato
RE
DAPI tdTomato
C
AAV-DIO-EGFP
SCN
Vip-ires-cre
D
SCN VIP Projection
DAPI EGFP
200 um
E
AAV-EGFP
or
AAV-Cre
RE
Vipr2-fl/fl
F
AP
-0.57
-0.87
G
Con
VIPR2 KO
Investigation index (F - NS %)
100
50
0
-50
CT12
CT18
CT12
CT18
H
Interaction (s)
300
150
0
CT12
CT18
CT12
CT18
I
Distance (cm)
3000
1500
0
CT12
CT18
CT12
CT18

**Fig. S7. Expression of VIPR2 in RE and influence of Vipr2 ablation in RE during familiarization session.**

(**A**) Illustration of Vipr2-tdTomato animal generation. (**B**) Representative images of tdTomato expression in RE. (**C**) Schematics for viral delivery. (**D**) Representative images of $SCN^{VIP}$ projections in RE. (**E**) Schematics for viral delivery. (**F**) Illustration of optic fiber placement. (**G** to **I**) Quantification of investigation index (G), total interaction (H), and total distance moved (I) during familiarization session of social priority assay with or without genetic ablation of Vipr2 in RE. n = 10, 11 for RE-EGFP and RE-Cre groups in (G to I). Two-way mixed ANOVA test followed by pairwise paired t-test and pairwise independent-sample t-test or non-parametric Two-way mixed ANOVA test followed by pairwise Wilcoxon Signed-Rank test and pairwise Wilcoxon Rank-Sum test was performed. Data are presented as mean ± SEM.

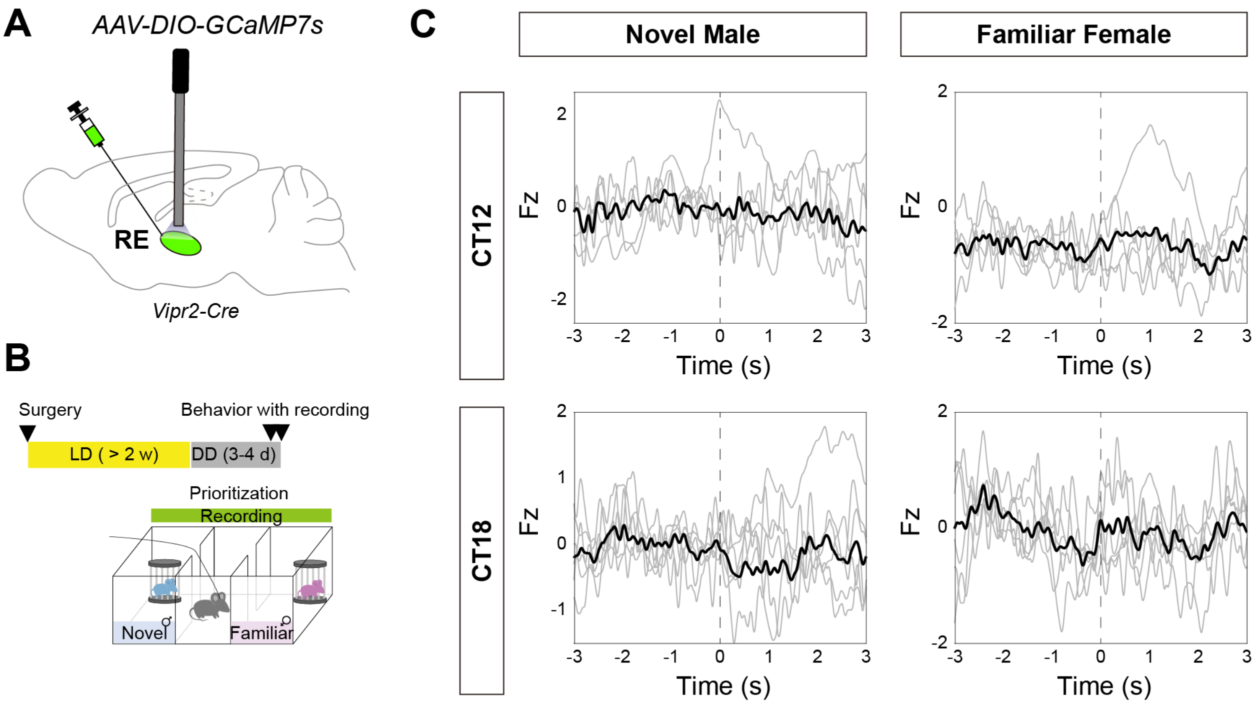
A
AAV-DIO-GCaMP7s
RE
Vipr2-Cre
B
Surgery
Behavior with recording
LD ( > 2 w)
DD (3-4 d)
Prioritization
Recording
Novel
Familiar
C
Novel Male
Familiar Female
CT12
CT18
Fz
Time (s)

**Fig. S8. In vivo calcium recording of $RE^{VIPR2}$ neuron during social priority.**

(**A**) Schematics for viral delivery. (**B**) Experimental scheme in vivo calcium recording during social priority assay. (**C**) PETH plot aligned to the entry of sniffing zone in each time point. n = 7 for $RE^{Vipr2}$-GCaMP7s (C). Each line represents calcium transients of individual animal (grey line) and an averaged calcium transient (black line). Dashed line indicates the onset of social interaction.

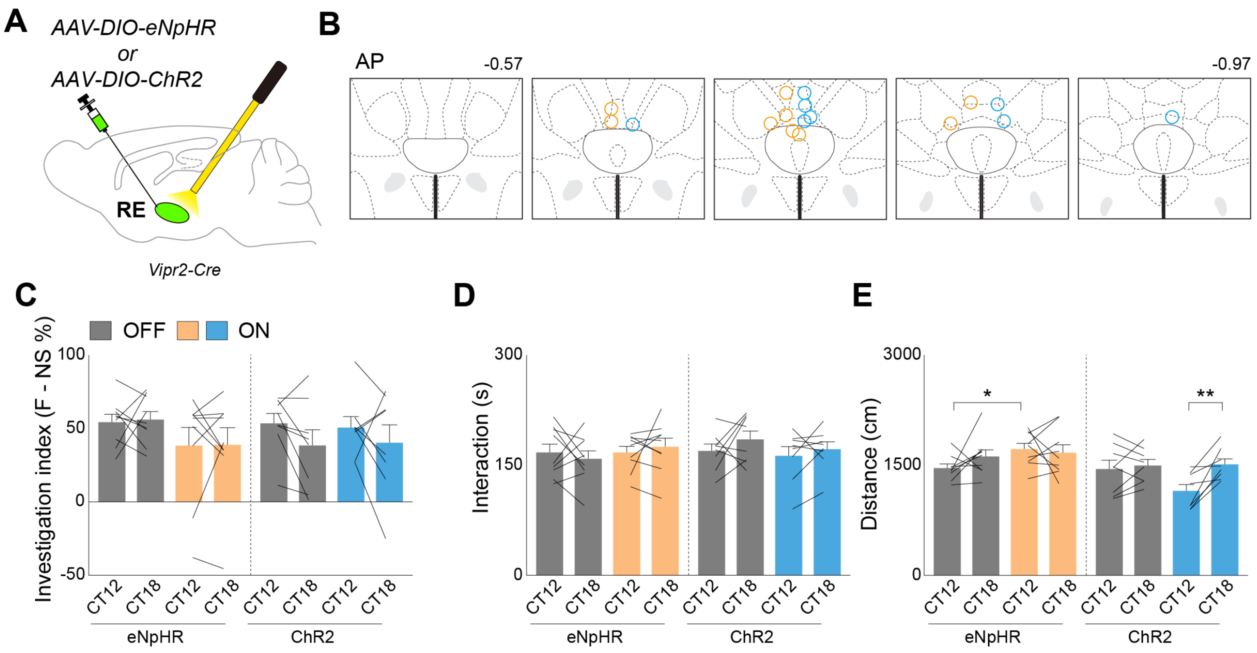

A
AAV-DIO-eNpHR
or
AAV-DIO-ChR2
RE
Vipr2-Cre
B
AP
-0.57
-0.97
C
OFF
ON
Investigation index (F - NS %)
100
50
0
-50
CT12
CT18
eNpHR
ChR2
D
300
150
0
Interaction (s)
E
3000
1500
0
Distance (cm)
*
**

**Fig. S9. Influence of optogenetic manipulation of $RE^{VIPR2}$ neurons during familiarization session.**

(**A**) Schematics for viral delivery. (**B**) Illustration of optic fiber placement. (**C** to **E**) Quantification of investigation index (C), total interaction (D), and total distance moved (E) during familiarization session of social priority assay with optogenetic inhibition in each time point. n = 9 for $RE^{VIPR2}$-eNpHR3.0 animals and n = 8 for $RE^{VIPR2}$-ChR2 animals in (C to E). For (C to E), RM Two-way ANOVA test followed by pairwise paired t-test or non-parametric Two-way ANOVA test was performed. Each line represents a repeated experiment within individual animal. Data are presented as mean ± SEM. $^{**}P < 0.01$; $^{*}P < 0.05$.

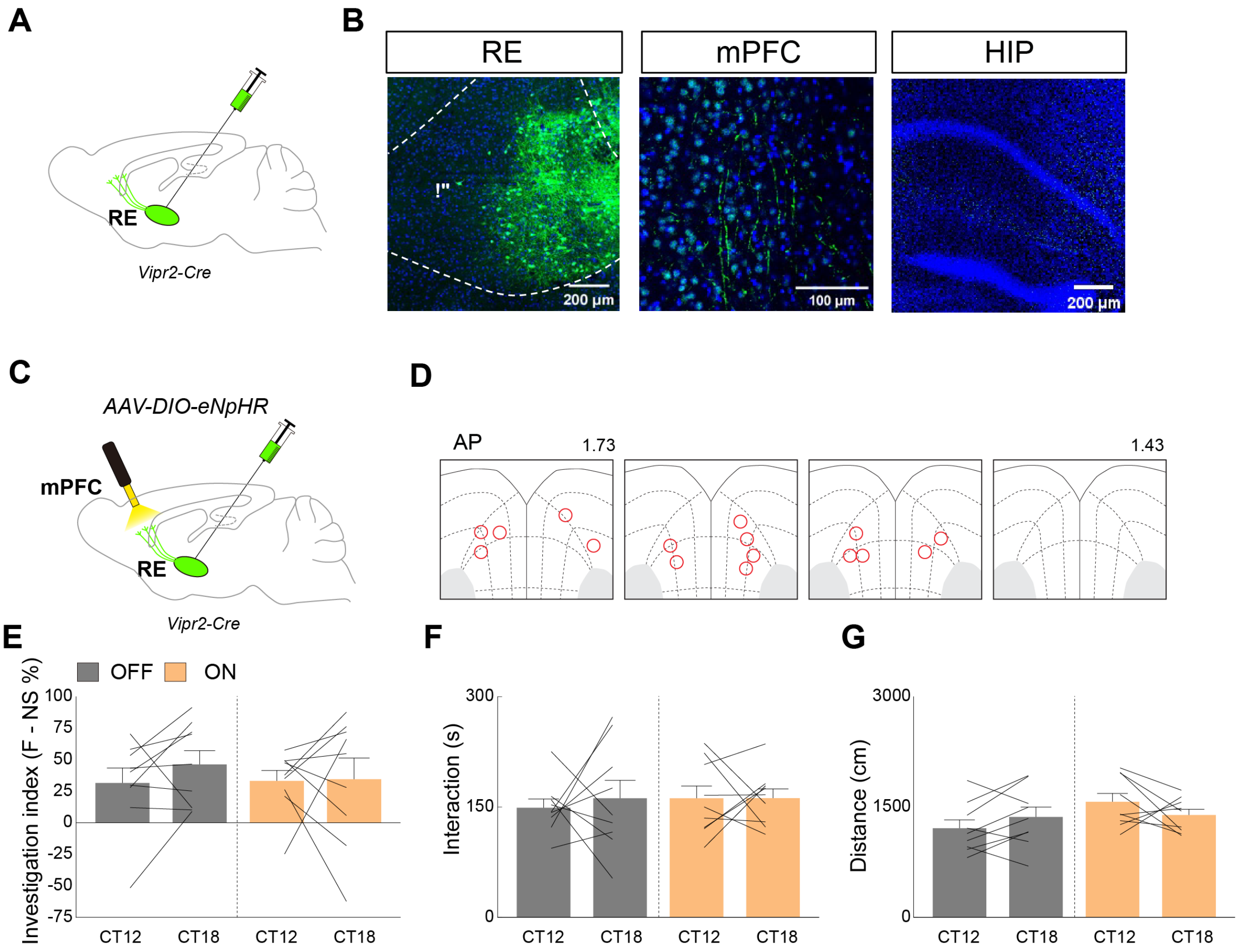
A
B
RE
mPFC
HIP
RE
Vipr2-Cre
200 μm
100 μm
200 μm
C
AAV-DIO-eNpHR
mPFC
RE
Vipr2-Cre
D
AP
1.73
1.43
E
OFF
ON
Investigation index (F - NS %)
100
75
50
25
0
-25
-50
-75
CT12
CT18
CT12
CT18
F
Interaction (s)
300
150
0
CT12
CT18
CT12
CT18
G
Distance (cm)
3000
1500
0
CT12
CT18
CT12
CT18

**Fig. S10. Neural projection of $RE^{Vipr2}$ neurons and influence of optogenetic inhibition of $RE^{VIPR2}$-mPFC circuit during familiarization session.**

(**A**) Schematics for viral delivery. (**B**) Fluorescence images of $RE^{Vipr2}$ neurons and their projection. (**C**) Schematics for viral delivery. (**D**) Illustration of optic fiber placement. (**E** to **G**) Quantification of investigation index (E), total interaction (F), and total distance moved (G) during familiarization session of social priority assay with optogenetic inhibition in each time point. n = 9 for $RE^{VIPR2}$-mPFC-eNpHR3.0 animals in (E to G). For (E to G), RM Two-way ANOVA test followed by pairwise paired t-test or non-parametric Two-way ANOVA test was performed. Each line represents a repeated experiment within individual animal. Data are presented as mean ± SEM.

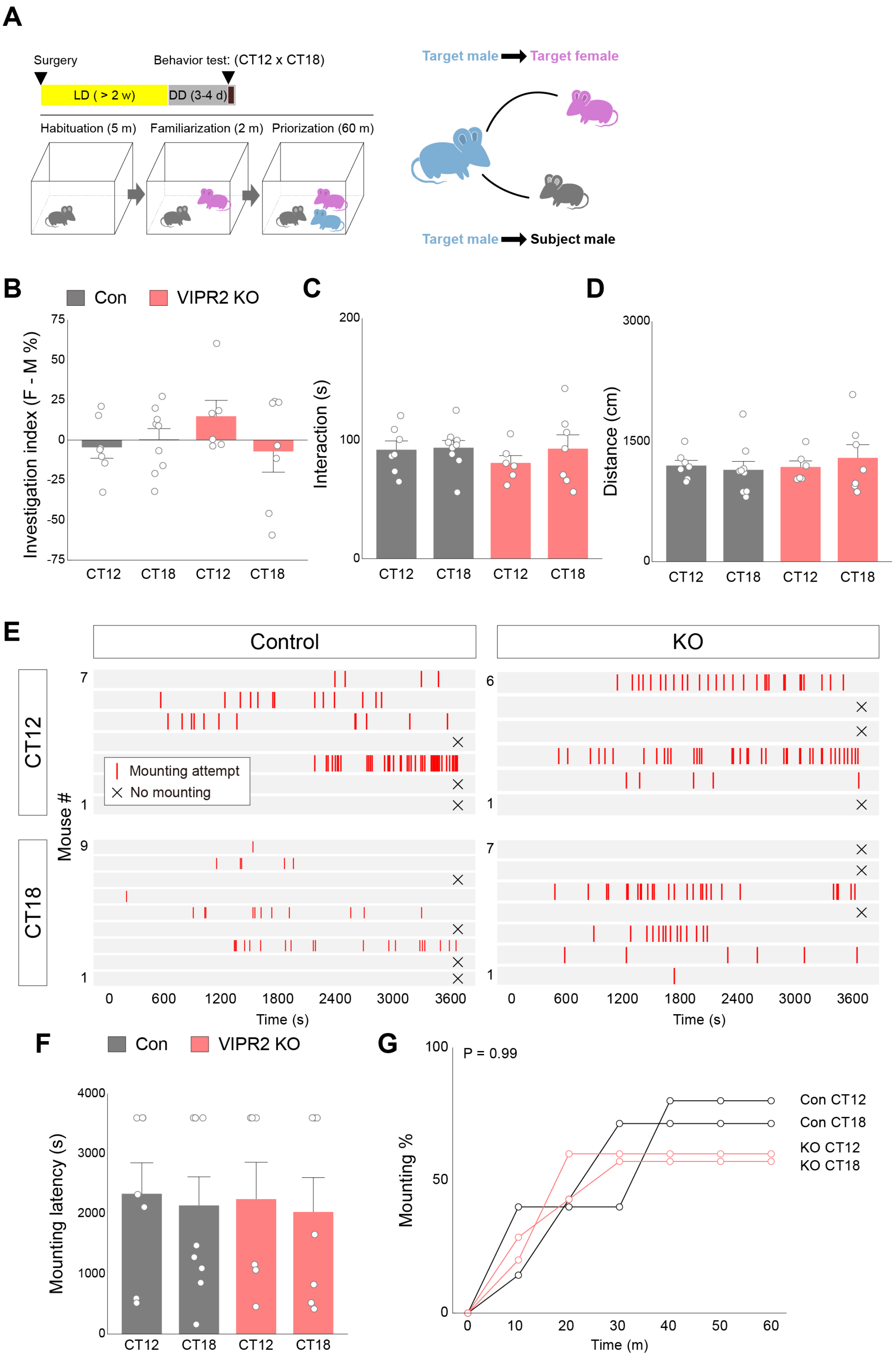
A
Surgery
Behavior test: (CT12 x CT18)
LD ( > 2 w)
DD (3-4 d)
Habituation (5 m)
Familiarization (2 m)
Priorization (60 m)
Target male
Target female
Target male
Subject male
B
Con
VIPR2 KO
Investigation index (F - M %)
CT12
CT18
CT12
CT18
C
Interaction (s)
D
Distance (cm)
E
Control
KO
CT12
CT18
Mouse #
Mounting attempt
No mounting
Time (s)
F
Con
VIPR2 KO
Mounting latency (s)
G
P = 0.99
Mounting %
Time (m)
Con CT12
Con CT18
KO CT12
KO CT18

**Fig. S11. Naturalistic group interaction and reproductive behavior by target male animals during TMI assay.**

(**A**) Experimental scheme of TMI assay (left) and illustration of group dynamics in TMI assay (right). (**B** to **D**) Quantification of investigation index (B), total interaction (C), and total distance moved (D) by target male mice during TMI assay. (**E**) Rastor plot of target male mice illustrating distribution of mounting attempts during TMI assay. (**F**) Latency to first mounting attempts by target male mice in each time point with or without genetic ablation of Vipr2 in RE. (**G**) Cumulative plot for the proportion of animals displaying mounting attempts in each time bin. n = 7, 9 for RE-EGFP CT12 and CT18 groups and n = 6, 8 for RE-Cre CT12 and CT18 groups. For (C to D) and (F), Two-way ANOVA test followed by pairwise independent-sample t-test or non-parametric Two-way ANOVA (ART) followed by pairwise Wilcoxon Rank-Sum test; for (G), Log-Rank test followed by pairwise Log-Rank test and Fisher's exact test followed by pairwise Fisher's exact test were performed. Each dot represents an individual animal. Data are presented as mean ± SEM.
